# Reducing Cofilin dosage promotes stress adaptation and thermotolerance in *Drosophila melanogaster* embryos

**DOI:** 10.1101/2025.01.02.631102

**Authors:** Natalie Biel, Subhashis Natua, Faizan Rashid, Divisha Biswas, Ting-Yu Wang, Tsui-Fen Chou, Auinash Kalsotra, Anna Marie Sokac

## Abstract

In addition to regulating actin dynamics, Cofilin also senses and responds to physiological stress and can determine cell survival outcomes. Yet, the full picture of Cofilin’s role in stress response is lacking. Here, we used imaging and RNA-seq methods to show that exposing early *Drosophila melanogaster* embryos to either acute or chronic heat stress (32°C) induces a Cofilin-mediated Actin Stress Response (ASR) and leads to upregulation of genes associated with both heat shock and Endoplasmic Reticulum (ER) unfolded protein responses. Reducing *cofilin* gene dosage (*cofilin^+/-^*) in heat-stressed embryos modulates all observed stress responses and partially rescues embryo survival. Unexpectedly we find that heat shock- and ER-stress response modulation arises because non-stressed *cofilin^+/-^* embryos already show upregulation of heat shock- and ER-stress response genes, prior to heat exposure. Our data support a model whereby *cofilin* heterozygosity activates specific stress responses associated with reduced protein homeostasis, thus priming embryos to be more resilient when they encounter subsequent heat stress. We conclude that Cofilin dosage serves as a determinant of stress outcomes, impacting both the actin cytoskeleton and inducible stress response pathways. In the embryo, we identify Cofilin as a novel link between inducible stress response and thermotolerance.

**Author summary:** Embryos regularly encounter physiological stresses that compromise their development and even put them at risk of death. These stresses can include hot or cold temperature shocks, chemical exposures, or metabolic imbalances carried over from the embryo’s parents. Here, we find a simple genetic manipulation that primes embryos to be more resilient to the stress of elevated temperature. By taking away one of the two normal copies of the gene *cofilin* in fruit flies, we generate embryos with improved survival upon exposure to heat stress. We find that the reduced dosage of Cofilin already introduces a low level of stress that then makes the embryos adapt so that they are more resilient when they encounter a subsequent stress. Our findings in fly embryos parallel previous observations showing that reducing *cofilin* gene copy number promotes the survival of neurons in mammals in both neurodegenerative and stroke contexts. However, our work is the first to consider Cofilin’s pro-survival activity using methods that capture the whole-system molecular response. As such, we reveal that Cofilin has a broad role to play in cellular stress response and makes an unanticipated contribution to stress adaptation and organismal thermotolerance.

## Introduction

Cofilin is best known as an essential regulator of actin cytoskeleton organization and dynamics (1). Cofilin binds actin filaments (F-actin) and drives their depolymerization or stabilization at low or high Cofilin-to-actin ratios, respectively (2, 3). In this way, Cofilin tunes the polymerization cycle to ultimately control the assembly and disassembly of actin-based structures in cells. Cofilin is itself regulated by upstream signaling that imparts instructions for how and when the actin cytoskeleton is remodeled (4). Conserved phosphatases, Slingshot and Chronophin, activate Cofilin by dephosphorylating its serine 3 residue. Conversely, kinases, including LIM kinase, inactivate Cofilin by phosphorylating the same residue. In some cases, Cofilin is also oxidized or subjected to other post-translational modifications to further modulate its localization or activity (4–6). In response to these signals, Cofilin then promotes F-actin remodeling that allows cells to carry out vital functions, such as migrating, dividing, or ingesting pathogens via phagocytosis (7).

In addition to F-actin regulation, Cofilin has several lesser recognized roles in cells. For example, Cofilin facilitates the import of actin monomers (G-actin) into the nucleus. Cofilin itself has a nuclear localization signal, and active dephosphorylated Cofilin can bind free G-actin as well as F-actin (8–12). Thus, Cofilin is thought to shuttle G-actin into the nucleus through its interaction with Importin-α9, though other mechanisms may exist (13–15). Once inside the nucleus, both Cofilin and actin influence gene expression through RNA Polymerase II (14, 16).

In another moonlighting role, Cofilin interacts with inducible stress responses in diverse ways. Cofilin’s normal regulation by oxidation positions it to serve as a redox sensor, and Cofilin becomes highly oxidized when the redox state of cells is perturbed (6, 17–19). If oxidative stress chronically persists, oxidized Cofilin moves to mitochondria, promoting swelling, downregulation of respiration and opening of permeability transition pores (6). Subsequent release of Cytochrome C into the cytosol then leads cells to commit to death programs including apoptosis, necrosis and ferroptosis (5, 6, 20). Thus, Cofilin can act as a potent initiator of mitochondrial-dependent cell death following persistent redox imbalance (5, 6, 20, 21).

Also related to stress functions, Cofilin is the principal driver of the inducible Actin Stress Response (ASR; also known as Cofilin-Actin Rod Formation) (22, 23). ASR is associated with several clinical conditions, including nemaline myopathy, ischemic brain and kidney injury, and neurodegenerative diseases such as Alzheimer’s and Huntington’s (24–30). ASR is induced by diverse stresses including oxidative stress, heat, hypoxia, ATP depletion, amyloid-β (Aβ) aggregation, and organismal aging (31–35). In the initial steps of ASR, Cofilin is hyperactivated, leading to the assembly of aberrant F-actin rods, in either the cytoplasm or nucleus, that are coated by high levels of dephosphorylated, and sometimes oxidized, Cofilin (11, 24, 32, 36–42). These rods become so stable that monomers within them do not turn over (32, 38, 43). In neurons, solid actin rods that form along axonal shafts physically block transport between the cell body and synapse, promoting synapse degeneration (34, 35, 44–46). In this context, Cofilin downregulation can inhibit rod assembly and be protective against cell dysfunction and death (22, 26, 47).

Thus, in playing its many roles, Cofilin has a complex relationship with cell health and survival. Under normal conditions, Cofilin activity is required for F-actin dynamics and cell function. However, in stress conditions, Cofilin can promote deleterious outcomes with its knockdown being protective against cell death. This shift in Cofilin function from beneficial to potentially harmful under physiological stress has generated interest in Cofilin as a target for clinical interventions (18, 26, 47, 48). Yet, the full extent to which Cofilin interacts with stress response and influences cell survival under stress is unclear. Regarding ASR, in particular, it remains debatable whether Cofilin’s activity in this stress response is protective or maladaptive for cells (32, 43, 49), and the function of Cofilin-actin rods is unknown. Furthermore, while rods serve as the main indicator of ASR, other rod-independent activities for Cofilin during this stress response have not been investigated.

We previously described an ASR that is mediated by Cofilin (*D.m.* Twinstar, Tsr) in *Drosophila* embryos reared under heat stress at 32°C (32, 50, 51). Embryos experiencing ASR show destabilization of cytoplasmic F-actin, prominent assembly of nuclear actin rods and diminished larval hatching rates. When we reduced the genetic dosage of *cofilin* to attenuate the ASR, we found that heat-stressed embryos were more likely to survive (32). In this current work, we sought to better understand how Cofilin alters embryo survival rates under heat stress. We initiated our study with the hypothesis that Cofilin negatively impacts embryo physiology upon heat stress due to its role in cytoplasmic F-actin disruption and rod formation. However, as we correlated survival data with quantitative analysis of these actin phenotypes for two distinct heat stress exposures, acute and chronic, we found that the observed actin disruptions were compatible with high rates of survival. To uncover how Cofilin’s activity during ASR may extend beyond actin organization, we used whole-embryo RNA sequencing (RNA-seq). By determining transcriptomic profiles of wild-type and *cofilin knockdown* embryos under non-stressed and heat-stressed conditions, we discovered a surprising interaction between Cofilin and the stress response. Specifically, we show that embryos with a reduced dose of *cofilin* (*cofilin^+/-^*) already exhibit signs of moderate heat shock and ER stress response, before encountering any elevation in temperature. We suggest that these embryos are primed by their genotype, such that when subsequently exposed to heat stress, they show improved survival.

## Results

### ASR is induced by both acute and chronic heat stress in embryos

We previously reported that wild-type *Drosophila* embryos reared under heat stress at 32°C exhibit disruption of cytoplasmic F-actin structures and the assembly of aberrant nuclear actin rods, indicative of a Cofilin-mediated ASR (32, 50, 51). In adult and aging mammalian cell types, the ASR can be induced by either acute or chronic physiological stress (22, 29, 30, 35, 48, 52, 53). To more systematically test whether the *Drosophila* embryo response is comparable to the mammalian response in this regard, we refined our prior protocol to include two distinct heat stresses: Specifically, embryos were reared with either transient “acute” or prolonged “chronic” heat stress at 32°C (1.5- or ≥ 12-hour exposure, respectively; Fig 1A). We considered the 32°C exposure a heat stress because *Drosophila* embryos can develop at 30-34°C but exhibit increasing defects and reduced survival (32, 54–59). In addition, while our conditions are not aligned with a typical “heat shock” exposure, which uses harsher temperatures of 37-42°C for a shorter duration (60–63), we nonetheless found by mass spectrometry analysis that heat shock proteins (HSPs) were upregulated at 32°C. From two independent experiments, we detected the induction of several small and large HSPs (Hsp22, 23, 26, 27, 60A, 68, 70Aa/Ab, 70Bb/Bc, 83 and Hsc70Cb/Hsp110; S1A Fig; S1 Table). Rearing at 18°C served as the control non-stressed condition because it falls within the range of temperatures that supports highest embryo survival rates, shows maximum canalization of gene expression between wild-type variant strains, and was identified in our earlier studies as optimal for robust morphogenesis (32, 50, 51, 59, 64).

**Fig 1.**
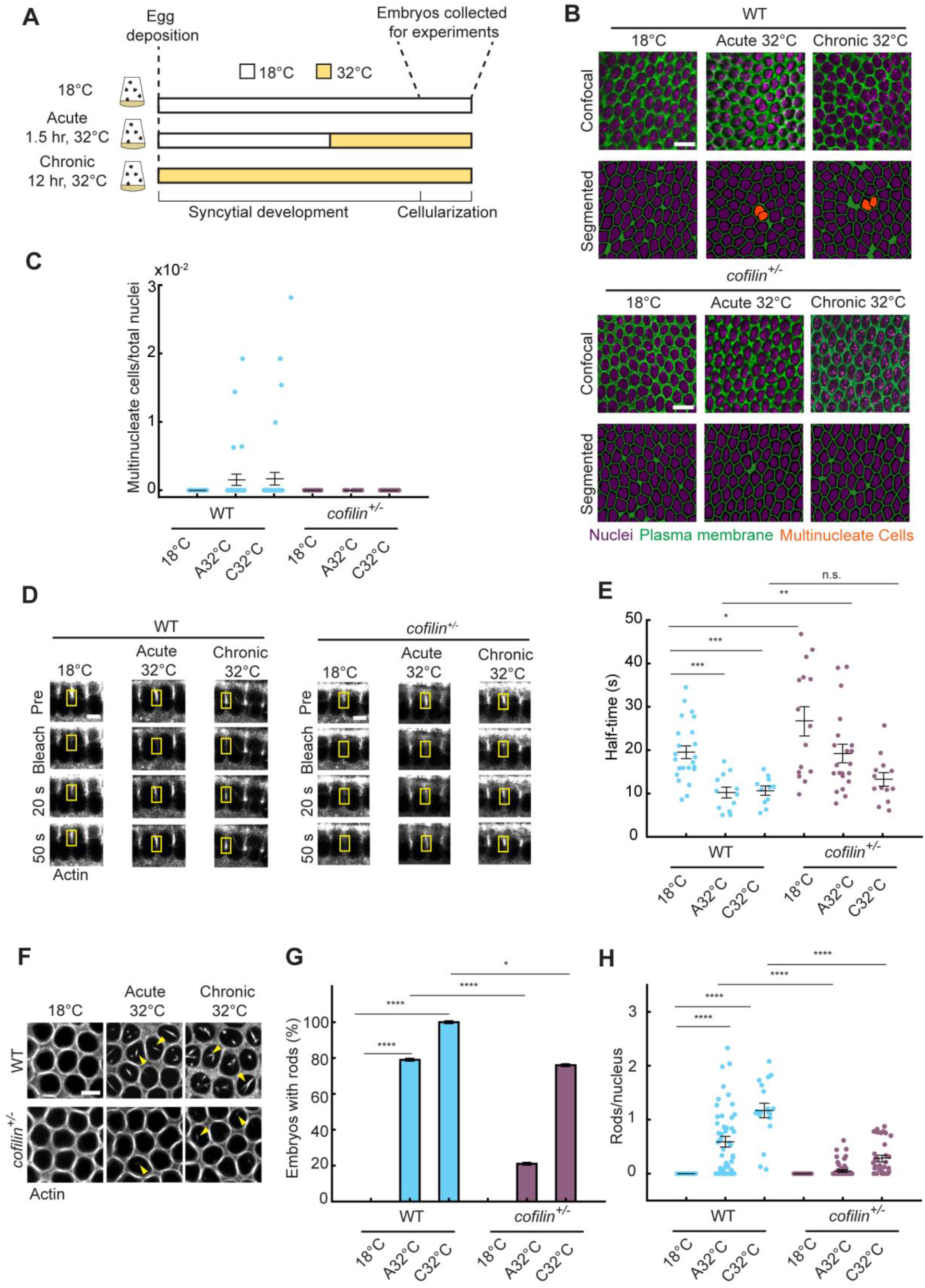
Cofilin drives the ASR in embryos exposed to either acute or chronic heat stress. (A) Schematic showing heat stress exposure times relative to developmental progression. Embryos collected at cellularization for all experiments. (B) Surface views and corresponding segmented images showing nuclei (Hoechst dye; purple) and plasma membrane furrows (Peanut; green) for WT and *cofilin^+/-^* embryos at cellularization at indicated conditions. Multinucleate cells highlighted by orange nuclei. Scale bar: 10 μm. (C) Dot plot showing ratio of multinucleate cells/total nuclei for WT (cyan) and *cofilin^+/-^* (purple) embryos at indicated conditions. Each point represents 1 embryo (n ≥ 15 total embryos per condition, with ≥ 70 cells assayed per embryo). This panel is treated as descriptive because differences between WT and *cofilin^+/-^*embryos for individual heat stress conditions are not statistically significant. (D) Cross-sections showing FRAP for furrow tip F-actin (G-actin^Red^) in WT and *cofilin^+/-^* embryos at indicated temperatures. Frame “pre” collected immediately prior to bleaching; frames 20 seconds (s) and 50s collected after bleaching. Yellow box indicates bleached region. Scale bar: 5 μm. (E) Dot plot showing half-time to recovery after photobleaching furrow tip F-actin for WT (cyan) and *cofilin^+/-^* (purple) embryos at indicated temperatures. Each point represents 1 embryo (n ≥ 11 embryos per condition, with 1 furrow bleached per embryo). (F) Surface views showing G-actin^Red^ incorporated into furrow tips and nuclear actin rods (yellow arrowheads) in WT and *cofilin^+/-^* embryos at indicated temperatures. Scale bar: 5 μm. (G) Bar plot showing percentage of WT (cyan) and *cofilin^+/-^* (purple) embryos with rods at indicated temperatures. (n ≥ 18 total embryos per condition, with ≥ 80 nuclei assayed per embryo). (H) Dot plot showing rods per nucleus for WT (cyan) and *cofilin^+/-^* (purple) embryos at indicated temperatures. Each point represents 1 embryo (n ≥ 18 embryos per condition, with ≥ 80 nuclei counted per embryo). *The mean, shown as horizontal lines in (C), (E) and (H) or bars in (G), was calculated from n = 3 independent biological replicates. Error bars represent standard error of the mean (SE)*. *p-values for (C), (E) and (H) calculated using Student’s t-test*. *p-values for (G) calculated using χ^2^ contingency analysis*. *\*p < 0.05, **p < 0.01, ***p < 0.001, ****p < 0.0001; p* ≥ *0.05 is not significant (n.s.)*.

Using these acute and chronic heat stress conditions, we first assessed the integrity of the cytoplasmic F-actin cytoskeleton. We focused on the first event in morphogenesis, which is cellularization of the syncytial embryo (65). Successful cellularization critically depends on stable F-actin structures forming at the tips of invaginating cleavage furrows and normally results in the generation of an epithelial sheet of hexagonally arrayed mononucleate cells (65). When F-actin at furrow tips is destabilized, some furrows regress, leaving multinucleate cells within the plane of the epithelial sheet (51, 66–68). To visualize multinucleation, we fixed wild-type embryos immediately after exposure to either acute or chronic heat stress at 32°C (Fig 1A), and stained using the furrow marker, Septin (*D.m.* Peanut, Pnut; Fig 1B). We found that the percentage of embryos showing multinucleation increased following either acute or chronic heat stress as compared to non-stressed controls at 18°C (13.7 ± 1.3% or 9.7 ± 0.3% multinucleate embryos, respectively; mean ± SE). This corresponded to an increase in the mean ratio of multinucleate cells to total nuclei in heat-stressed embryos (Fig 1C).

Because multinucleation can be caused by reduced F-actin stability at furrow tips, we used fluorescence recovery after photobleaching (FRAP) to assay F-actin turnover in live heat-stressed embryos (32, 69, 70). FRAP measures the rate of actin monomer flux within filaments, where faster recovery indicates that filaments are less stable (71). To visualize furrow tip F-actin, we injected non- or heat-stressed embryos with rhodamine-conjugated non-muscle G-actin (G-actin^Red^) at 30 minutes post egg deposition. G-actin^Red^ incorporates into F-actin structures and preserves normal F-actin function in developing embryos (32, 69, 70). Non-stressed embryos were imaged at 18°C, while heat-stressed embryos were imaged on a temperature-controlled stage at 32°C. We found that the half-time to recovery was significantly faster for furrow tip structures upon either acute or chronic heat stress (Figs 1D and 1E), though the percent mobile fraction did not change relative to the 18°C control condition (S1B Fig). Thus, the dynamic population of furrow tip F-actin is destabilized in wild-type embryos challenged by heat stress of both durations. To next test for disruption of the nuclear actin cytoskeleton, we examined the assembly of nuclear actin rods in heat-stressed embryos. These rods are composed of parallel bundles of highly twisted F-actin and indicate induction of the ASR (36–38). To visualize rods, we injected non- or heat-stressed wild-type embryos with G-actin^Red^, followed by live imaging at 18°C or 32°C, respectively (32, 50). We collected single plane images that bisect nuclei. At this plane, F-actin is visible in furrow structures that encircle the dark nuclei (Fig 1F). Inside the nuclei, actin rods were prominent in wild-type embryos exposed to either acute or chronic heat stress but were completely absent in non-stressed embryos (Fig 1F). The fraction of wild-type embryos with rods and the number of rods per nucleus were most elevated in the chronic heat stress condition (Figs 1G and 1H). In addition, actin rods were larger in diameter in wild-type embryos exposed to chronic versus acute stress (S1C Fig). Nuclear actin rods in both heat stress conditions were “stable” as determined by a lack of recovery of G-actin^Red^ signal within rods following FRAP (S1D and S1E Figs). This rod stability is consistent with observations made for rods in mammalian ASR (38, 43). In summary, ASR in *Drosophila* embryos resembles the mammalian ASR in being inducible by either acute or chronic heat stress. In embryos, cytoplasmic F-actin destabilization occurs simultaneously with the formation of actin rods in the nucleus, and the number and size of rods increases with the severity of heat stress.

### Embryo ASR is Cofilin-dependent

Cofilin is the main driver of the mammalian ASR in both acute and chronic stress conditions (22, 23, 30, 35, 48, 49, 53). We previously showed that the ASR in heat-stressed embryos is Cofilin-dependent by reducing the maternal dose of the *cofilin* gene (*i.e.* embryos were harvested from mothers heterozygous for *tsr^1^*, a hypomorphic allele of *cofilin*) (32). However, at that time, we did not systematically test for Cofilin’s involvement in ASR in embryos subjected to both acute and chronic heat stress.

To now examine Cofilin’s role in the ASR after each heat stress exposure, we employed our maternal heterozygosity approach again and validated that knockdown in mothers reduces the Cofilin protein level to 62.5 ± 12.4 % of the wild-type level in embryos (mean ± SE; S1F Fig). For the cytoplasmic F-actin cytoskeleton, we saw no multinucleation in these *cofilin*^+/-^ embryos following either acute or chronic heat stress at 32°C (Fig 1B). Accordingly, the mean ratio of multinucleate cells to total nuclei equaled zero in all heat-stressed *cofilin*^+/-^ embryos (n ≥ 15 total embryos per condition, with ≥ 70 cells assayed per embryo; Fig 1C). Thus, *cofilin* knockdown completely rescued furrow regression in heat-stressed embryos.

To ask if furrow regression is prevented because F-actin remains stable in *cofilin*^+/-^ embryos, we assayed the turnover rates of furrow tip F-actin using FRAP. Non-stressed *cofilin*^+/-^ embryos at 18°C showed a slower half-time to recovery compared to non-stressed wild-type embryos (Figs 1D and 1E). Slower recovery indicates increased F-actin stability, which is predicted when Cofilin’s F-actin depolymerization activity is decreased by reduced Cofilin dosage (1–3). Similarly, after acute heat stress, *cofilin*^+/-^ embryos showed slower recovery than their wild-type counterparts (compare 18.8 ± 2.0 and 10.2 ± 1.2 seconds, respectively; mean ± SE; Fig 1E), suggesting *cofilin* knockdown countered cytoplasmic F-actin destabilization under this condition. For chronic heat stress, *cofilin*^+/-^ embryos also showed a slower average half-time to recovery compared to wild-type embryos under the same condition, though not significantly so (compare 13.4 ± 1.5 seconds and 10.9 ± 1.0 seconds, respectively; mean ± SE; Fig 1E). The percent mobile fraction was constant across conditions for *cofilin*^+/-^ embryos, suggesting that dynamic F-actin population did not change (S1B Fig). These results show that *cofilin* knockdown reduces but does not entirely alleviate disruption of the cytoplasmic F-actin cytoskeleton in heat-stressed embryos, particularly in the context of chronic heat stress when additional factors, such as oxidation of actin or actin binding proteins, may influence actin turnover (72, 73).

To then test for assembly of nuclear actin rods as the signature read-out of ASR in *cofilin*^+/-^ embryos, we used the same G-actin^Red^ injection and imaging method as described above. In 18°C non-stressed *cofilin*^+/-^ embryos, rods were absent inside nuclei, whereas nuclear rods were seen in *cofilin*^+/-^ embryos exposed to acute or chronic heat stress (Fig 1F). However, we found that the fraction of *cofilin*^+/-^ embryos with rods and the number of rods per nucleus were significantly decreased compared to wild-type embryos, for both stress conditions (Figs 1G and 1H). Nuclear actin rods were also thinner in *cofilin*^+/-^ embryos for both stress conditions (S1C Fig). Despite their reduced size, rods in *cofilin*^+/-^ embryos were stable as determined by photobleaching (S1D and S1E Figs), suggesting that a lack of turnover of actin monomers is a general feature of rods in embryos, regardless of stress condition or genotype. In summary, these results show that Cofilin supports the formation of actin rods in embryonic ASR upon both acute and chronic heat stress exposure, analogous to mammalian ASR.

### Rod formation is associated with reduced survival, but is not its sole determinant

It remains unclear what benefit the ASR imparts to stressed cells. Is it fundamentally protective or maladaptive? Mammalian ASR is suggested to be either protective or pathological in acute or chronic stress conditions, respectively (34, 35, 43–46, 49). Previously, we described the ASR in embryos as maladaptive, because *cofilin*^+/-^ embryos, with diminished ASR, survived to larval hatching more frequently than wild-type embryos with an intact ASR (32). However, we only tested the relationship between ASR and survival for chronically stressed embryos.

To more thoroughly evaluate if ASR can be protective in embryos, we revised our hatching experiment to include both acute and chronic heat stress exposures. In addition, we controlled for the genetic background of the “wild-type” embryos to avoid confounding survival effects that might arise from outcrossing the *cofilin* mutant stock. Specifically, the *cofilin*^+/-^ mothers are prepared by outcrossing parental wild-type Oregon-R (Ore-R) males with females from a stock carrying the balanced *tsr^1^* allele of *cofilin*. In general, outcrossing alone improves survival by diluting undesirable mutations that accumulate over time in inbred lab stocks (74–76), which would then misleadingly inflate the survival rate of *cofilin*^+/-^ embryos when compared with inbred wild-type embryos. To avoid this in our control wild-type embryos, we now outcrossed Ore-R males to females homozygous for *white^1118^*, which is a marker mutation commonly used in genetic experiments as a control (77). Thus, for all subsequent experiments both the wild-type and *cofilin*^+/-^ embryos are generated via an outcrossing strategy.

We performed larval hatching assays on a total of 600 embryos from three independent experiments for each non- and heat-stressed condition. Embryos exposed to either acute or chronic heat stress were hand-selected at cellularization and shifted to 22°C to continue development to larval hatching (Fig 1A). Control non-stressed embryos were reared at 18°C until cellularization and then shifted to 22°C. We found that control wild-type embryos exposed to either acute or chronic heat stress prior to cellularization show significantly reduced hatching as larvae, with reduced survival most pronounced for chronic heat stress (Fig 2A; S2A Fig). These results are consistent with other reports that embryos exposed to temperatures ≥ 30°C, even for short times, show reduced viability (32, 54–59).

**Fig 2.**
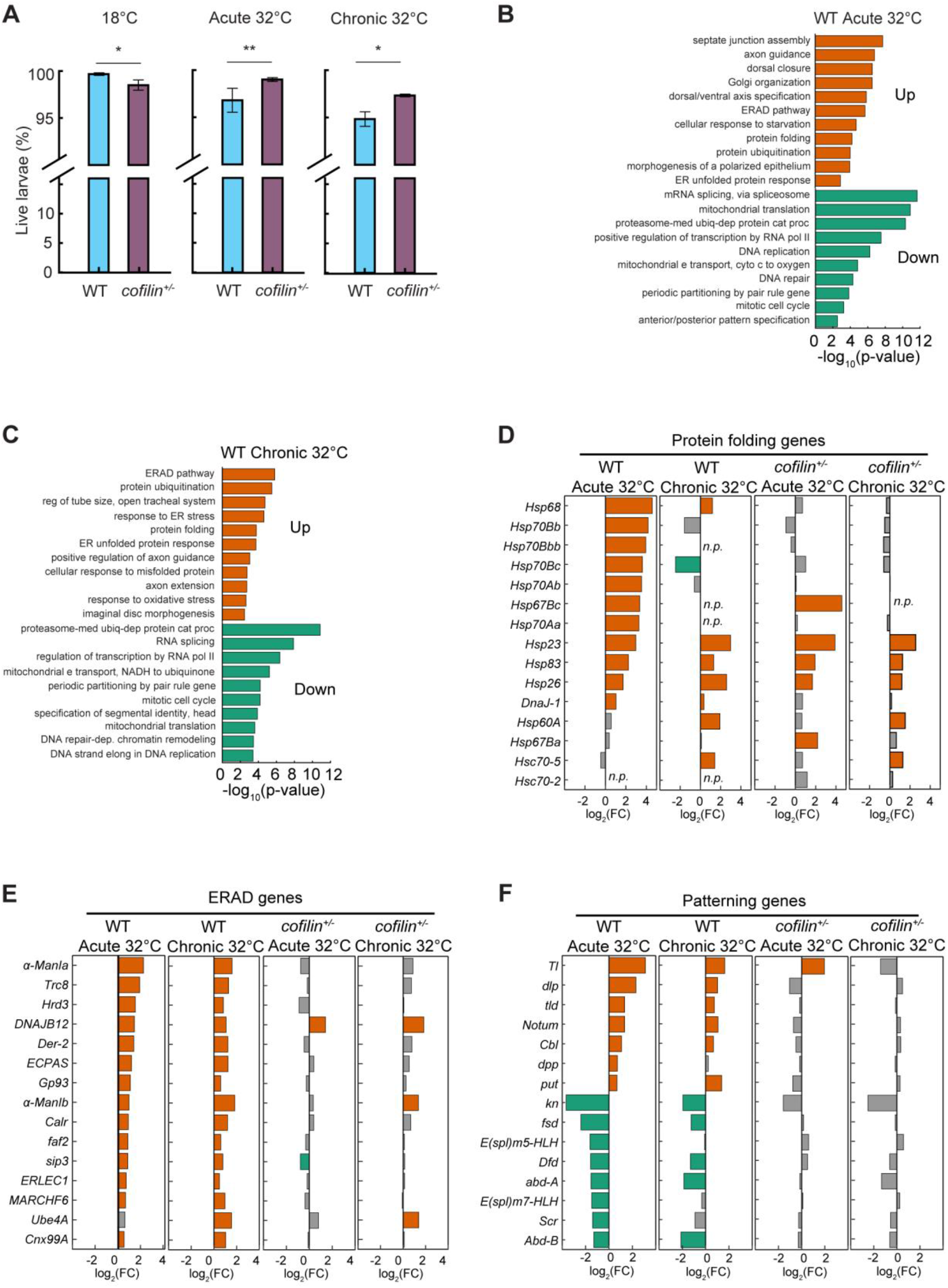
Heat-stressed wild-type embryos show reduced survival and upregulation of HSPs and ER unfolded protein response genes. (A) Bar plots showing percentage of survival for larvae for wild-type (WT; cyan) and *cofilin^+/-^* (purple) embryos at indicated conditions. (B, C) Bar plots showing -log_10_(p-value) for selected highly significant GO Biological Processes for upregulated (orange) or downregulated (green) gene sets for acute 32°C (B) or chronic 32°C (C) heat-stressed WT embryos versus 18°C non-stressed WT embryos. Abbreviated processes are: proteasome-mediated ubiquitin-dependent protein catabolic process; mitochondrial electron transport, cytochrome c to oxygen; regulation of tube size, open tracheal system; mitochondrial electron transport, NADH to ubiquinone; DNA repair-dependent chromatin remodeling. (D, E and F). Bar plots showing log_2_(FC) values for upregulated (orange) or downregulated (green) genes from highly significant ontologies related to: Protein folding (D); ERAD pathway (E); and Patterning (F) for WT or *cofilin^+/-^* embryos at indicated conditions. For Patterning, genes in the top 7 bars or bottom 8 bars are associated with dorsal/ventral or anterior/posterior patterning, respectively. Genes with no significant change in expression shown in gray. Abbreviation in (D) is: not present (n.p.). *The mean, shown as bars in (A), was calculated from n = 600 embryos total for each condition from 3 independent replicates of 200 embryos per replicate. Error bars represent standard error of the mean (SE). p-values calculated using χ^2^ contingency analysis. *p < 0.05, **p < 0.01. Differential gene expression data in (B-F) were based on n = 6 biological replicates for 18°C WT and acute 32°C WT or n = 5 for chronic 32°C WT; and n = 6 for all conditions for cofilin^+/-^. Significance value cut-off for differentially expressed genes was FDR < 0.05; and significant ontologies p < 0.01*.

We then examined the survival of *cofilin*^+/-^ embryos. At 18°C with no heat stress, these embryos had slightly lower survival rates than their wild-type counterparts (Fig 2A; S2A Fig). However, following either acute or chronic heat stress, the *cofilin*^+/-^ embryos showed a small but significant survival advantage over wild-type embryos (Fig 2A; S2A Fig). Thus, reducing the dosage of Cofilin makes embryos more resilient, somehow countering the negative effects of heat stress on embryo survival.

These results argue that the Cofilin-dependent ASR is maladaptive in both acute and chronic heat stress scenarios. However, some aspects of our data fail to align with the simple explanation that the actin phenotypes we observe during the ASR are a one-to-one predictor of embryo survival. For example, *cofilin*^+/-^ embryos exposed to chronic heat stress showed the same extent of cytoplasmic F-actin destabilization as wild-type embryos but had a higher survival rate (Figs 1E and 2A). Also, in the chronic stress condition, 76.14 ± 0.05% of *cofilin*^+/-^ embryos assembled nuclear actin rods (mean ± SE), but fewer than 3% of these embryos failed to hatch (Figs 1G and 2A). So, while we find that the severity of actin changes correlates with decreased survival, our data do not support that cytoplasmic F-actin destabilization or rod formation serve as the exclusive drivers of reduced survival in wild-type embryos. Instead, we suggest that ASR and actin changes act together with other unknown responses to compromise survival in heat-stressed wild-type embryos, and that *cofilin* knockdown mitigates these responses, making embryos more resilient to the stress.

### In wild-type embryos heat stress induces gene expression consistent with heat and ER stress responses

To identify responses that accompany the ASR and actin phenotypes in heat-stressed embryos, we took an unbiased approach and performed whole-embryo RNA-seq. We considered whole-embryo sequencing reasonable at this early stage of development because cell-type specification is only beginning. Outcrossed wild-type embryos were exposed to acute or chronic stress at 32°C as in the earlier experiments (Fig 1A), and poly-A RNA was isolated at Bownes Stage 5, corresponding to early/mid cellularization (78). Six independent biological replicates were analyzed for the 18°C non-stressed and acute 32°C heat-stressed wild-type conditions, and five biological replicates were analyzed for the chronic 32°C heat-stressed wild-type condition. Following sequencing, we identified significantly up- or down-regulated genes in heat-stressed wild-type embryos compared to non-stressed siblings at 18°C (FDR < 0.05). Because the maternal-to-zygotic transition (MZT) is ongoing at Stage 5, detected genes can represent both maternally and zygotically contributed transcripts (79). To validate RNA-seq results, we compared them with expression profiles determined by quantitative PCR (qPCR). The same differential expression trends were observed by both methods for 13 out of 14 cases (>92% reproducibility; S2B Fig; S2 Table). Finally, we performed gene enrichment analysis on each set of up- or down-regulated genes per condition using PANGEA to identify the most statistically significant biological processes (p < 0.01) (80). Complete gene sets and results from enrichment analysis are provided in S3 and S4 Tables.

For wild-type embryos exposed to acute heat stress, 2052 genes were significantly upregulated and 1931 downregulated (FDR < 0.05; S2C Fig; S3 Table). A response to heat stress was evident by increased expression of protein folding genes, including HSPs (Figs 2B and 2D; S4 Table) (81). Induction of HSPs suggests compromised protein homeostasis (i.e. proteostasis) upon heat exposure; and this interpretation was reinforced by coincident upregulation of components of the Endoplasmic Reticulum (ER) unfolded protein response and the ER Associated Degradation (ERAD) pathway, which helps to clear unfolded proteins from the ER (Figs 2B and 2E; S2E Fig; S4 Table) (82, 83). In addition, genes involved in cell adhesion, signaling, cellular transport, metabolism and cytoskeletal remodeling were also upregulated consistent with transcriptomes previously reported for heat-stressed yeast and mammalian cells (S4 Table) (84, 85). Among the downregulated genes, there was apparent repression of processes related to transcription, splicing, DNA repair, and mitochondrial activity, also consistent with known cellular reactions to heat stress and consequent protein unfolding (Fig 2B; S2C and S2F Figs; S4 Table) (81, 86–92). Together, the signatures of upregulated and downregulated genes suggests that embryos exposed to acute heat stress are responding to a loss in proteostasis.

Developmental genes are both upregulated and downregulated in these acutely stressed embryos (Figs 2B and 2F; S4 Table). Normally at stage 5, embryos are undergoing extensive transcriptional remodeling during the MZT, the body plan is being established, and morphogenesis is starting (79, 93–97). We observed that for the acutely stressed embryos, genes involved in dorsal/ventral patterning tend to be upregulated while genes involved in anterior/posterior patterning tend to be downregulated (Figs 2B and 2F; S4 Table). Among the upregulated genes, there is also a striking representation of cellular machinery relevant to building epithelial and neuronal tissues. For example, biological processes include morphogenesis of a polarized epithelium and axon guidance (Fig 2B; S4 Table). So, besides induction of well-established inducible stress responses, embryos also modify their developmental programs. Based on the relatively high survival rate determined for these embryos, this developmental response is not considered to be overtly perturbative. However, it remains unclear if these modifications to developmental gene expression are simply a consequence of heat stress that embryos are robust against, or if the modifications represent an effort to resist specific challenges to tissue building and patterning at elevated temperatures.

For wild-type embryos exposed to chronic heat stress, a similar number of genes were differentially expressed as in the acute condition, with 1967 upregulated and 1654 downregulated (FDR < 0.05; S2D Fig; S3 Table). What’s more, many of the significant biological processes overlapped between the chronic versus acute stress conditions (Figs 2B and 2C; S4 Table). Again, HSPs and genes associated with ER unfolded protein response and ERAD were upregulated, while transcription, splicing and DNA repair genes were downregulated (Figs 2D and 2E; S2D-S2F Figs; S4 Table). So, as observed in acute heat stress, chronically heat-stressed embryos also show signs of reduced proteostasis. In addition, some unique indicators of stress emerged following chronic heat exposure, such as the upregulation of oxidative stress genes, consistent with the more severe stress condition increasingly compromising embryo fitness (Fig 2C; S4 Table). However, we noted that the fold change (FC) among upregulated HSPs was frequently lower in the chronic than acute condition (Fig 2D), perhaps because these embryos are exhibiting adaptation to heat as a prolonged stressor rather than a transient assault (for example, compare log_2_(FC) = 1.1 or 4.5 for Hsp68 after chronic or acute stress, respectively, and Hsp70Ab was absent in the chronic stress condition; S3 Table).

Considering development, processes related to epithelial and neuronal tissue building remained significant for the upregulated genes in chronically heat-stressed embryos (Fig 2C; S4 Table). Also, anterior/posterior patterning genes were downregulated in the chronic stress condition (Figs 2C and 2F; S4 Table). Somewhat unique in comparison to the acute heat stress condition, dorsal/ventral patterning was no longer represented among the most significant biological processes for the upregulated genes in chronically heat stressed embryos, although genes relevant to dorsal/ventral patterning were still upregulated (Fig 2F; S4 Table).

Overall, our data for wild-type embryos show that both acute and chronic heat stress at 32°C activate transcriptional responses in embryos in addition to the ASR and observed actin phenotypes. Specifically, heat shock and ER unfolded protein responses are induced (Figs 2B-2E; S2E Fig; S4 Table). Alongside these known stress pathways, changes in developmental gene expression accompany the ASR (Figs 2B, 2C and 2F; S4 Table). The extent of differential gene expression for both stress response and development can vary with severity of stress, perhaps depending on when the embryo first encounters the stress (i.e. as an embryo for acute stress versus as an oocyte for chronic stress). We conclude that heat-stressed embryos have multiple coincident responses that extend beyond the known ASR and actin cytoskeleton perturbations. These combined stress responses, however, do not fully protect against elevated temperature, as judged by reduced embryo survival (Fig 2A; S2A Fig).

### Within-genotype comparisons suggest *cofilin knockdown* embryos mount a muted transcriptional response to heat stress

Given that cofilin knockdown leads to improved survival compared to the wild-type genotype (Fig 2A; S2A Fig), and that this improved survival may not only depend on reducing the degree of actin disruption and rod formation, we wanted to also determine the transcriptional response of *cofilin*^+/-^ embryos to heat stress. We repeated RNA-seq as described above, performing six independent biological replicates per stress condition for *cofilin*^+/-^ embryos. In both this run of RNA-seq and the prior with the outcrossed wild-type embryos, two replicates of an independent strain (inbred Ore-R) were included per condition to facilitate assessment of sequencing-run-associated variation between the two datasets. Significantly up- or down-regulated genes (FDR < 0.05) were identified by comparing heat-stressed *cofilin*^+/-^ embryos to their non-stressed siblings reared at 18°C. Enrichment analysis was performed on each differentially expressed gene sets per condition, using PANGEA to identify the most highly significant biological processes (p < 0.01) (80). Complete gene sets and results from enrichment analysis are provided in S3 and S4 Tables.

After acute heat stress at 32°C, 344 genes were upregulated and 273 downregulated in *cofilin*^+/-^ embryos (FDR < 0.05; S3A Fig; S3 Table). This was a drastically muted response compared to that seen in the acute stress condition for wild-type embryos (compare a total 617 of differentially expressed genes in *cofilin*^+/-^ versus 3983 genes in wild-type; S2C and S3A Figs). As in wild-type, some HSPs and genes associated with response to heat were upregulated in stressed *cofilin*^+/-^ embryos (Fig 2D; S3C Fig; S4 Table). However, upregulated HSP genes were fewer compared to wild-type embryos; and no Hsp70s nor Hsp68 were upregulated in *cofilin*^+/-^ embryos (Fig 2D; S3 Table). Also notable, ER unfolded protein response and ERAD were absent from the biological processes identified for the upregulated genes in *cofilin*^+/-^ embryos (Fig 2E; S2E and S3C Figs; S4 Table). Furthermore, the large-scale downregulation of genes associated with transcription, splicing, DNA repair, and mitochondrial processes which emerged as a major part of the response to acute heat stress in wild-type embryos, was milder for the *cofilin*^+/-^ embryos (S2F and S3C Figs; S4 Table). Combined, these results suggest that *cofilin*^+/-^ embryos detect acute heat stress at 32°C, but display a diminished response compared to wild-type counterparts.

Even when challenged by chronic heat stress, *cofilin*^+/-^ embryos showed a weaker response than wild-type embryos, where 820 genes were significantly upregulated and 716 downregulated (compare a total of 1536 differentially expressed genes in *cofilin*^+/-^ versus 3621 genes in wild-type; FDR < 0.05; S3B Fig; S3 Table). Among the upregulated genes in the chronically heat-stressed *cofilin*^+/-^ embryos, HSPs were observed (Fig 2D), but the number of HSPs identified as differentially expressed was less than in wild-type under the same condition. Biological processes related to ER unfolded protein stress response and ERAD were not significantly represented for the *cofilin*^+/-^ embryos (Fig 2E; S2E and S3D Figs; S4 Table). In contrast to wild-type embryos, genes involved in transcription and DNA repair did not show strong downregulation (S2F and S3D Figs; S4 Table). Thus, *cofilin*^+/-^ embryos consistently showed fewer transcriptomic changes between the non-stressed and heat-stressed conditions than wild-type embryos. While their response to heat appeared diminished, the *cofilin*^+/-^ embryos did show downregulation of genes involved in cytoplasmic translation and ribosome assembly (S3D Fig; S4 Table), suggesting that proteostasis may be uniquely managed in these embryos (98, 99). Combined, our observations for *cofilin*^+/-^ embryos show that their heat stress responses, including both observable actin phenotypes as well as transcriptional changes, deviate from the wild-type response in a manner that also corresponds with improved survival.

### Between-genotype comparisons reveal a distinct, but robust transcriptional stress response in *cofilin* knockdown embryos

To more directly determine how heat stress response differs in *cofilin*^+/-^ embryos compared to wild type, we modified our analysis: Rather than look at the “within-genotype” response to heat stress (i.e. wild-type 32°C versus wild-type 18°C, or *cofilin*^+/-^ 32°C versus *cofilin*^+/-^ 18°C), we compared “between-genotypes”. Specifically, we performed pairwise comparisons between the differentially expressed genes for *cofilin*^+/-^ versus wild-type embryos for each heat stress condition (i.e. *cofilin*^+/-^ 32°C versus wild-type 32°C).

For acute heat stress response, we found that 2109 genes were upregulated, and 1413 genes were downregulated for *cofilin*^+/-^ versus wild-type embryos (FDR < 0.05; S3E Fig; S3 Table). Due to the prior within-genotype analysis, we were surprised to see that a subset of HSPs and their homologous heat shock cognate (Hsc) genes were upregulated in the *cofilin*^+/-^ response compared to wild type (Hsps 23, 67Ba, 67Bc, 83 and Hscs 70-2, 70-5, 70Cb; Fig 3A; S3 Table). That is, these embryos expressed heat shock chaperones at levels matching or possibly exceeding their wild-type counterparts, even though the within-genotype comparison made the *cofilin*^+/-^ response appear dampened. Interestingly, the ERAD pathway was identified by enrichment analysis of the downregulated genes, rather than the upregulated genes, suggesting that this pathway is not activated during the *cofilin*^+/-^ response to acute heat stress (Figs 3B and 3D; S4 Table). Similarly, dorsal/ventral patterning was associated with downregulated genes in the *cofilin*^+/-^ response (Figs 3C and 3D; S4 Table), whereas it had been associated with upregulated genes in the wild-type response. Processes related to transcription, DNA repair and anterior/posterior patterning were associated with the upregulated genes in the *cofilin*^+/-^ response, in contrast to their association with downregulated genes in the wild-type response (Figs 3C and 3D; S4 Table). So, the *cofilin*^+/-^ response to acute heat stress is not so much diminished as it is distinct from the wild-type response, with elevated levels of some chaperones present and other signatures of stress response modified. For chronic heat stress, we found that 694 genes were upregulated, and 297 genes were downregulated in *cofilin*^+/-^ versus wild-type embryos (FDR < 0.05; S3F Fig; S3 Table). This suggested less divergence between the *cofilin*^+/-^ and wild-type responses for this heat stress condition. Notably, though, all five of the Hsp70s were upregulated in the *cofilin*^+/-^ response compared to wild type, as well as Hsps23, 68 and 83 and Hsc70-2 (Fig 3A), showing that, as for acute heat stress, *cofilin*^+/-^ embryos express chaperones at levels at least commensurate with wild-type embryos. ERAD pathway and ER unfolded protein response were not identified in association with either up- or down-regulated genes, indicating that the *cofilin*^+/-^ and wild-type ER responses to chronic heat stress are comparable (Figs 3B and 3E; S4 Table). This was also largely the case for processes related to patterning (Figs 3C and 3E; S4 Table). So, again, and in contrast to the impression from the within-genotype comparisons, *cofilin*^+/-^ embryos retain a response to chronic heat stress; and, in fact, this response may not drastically deviate from the wild-type response to chronic heat stress.

**Fig 3.**
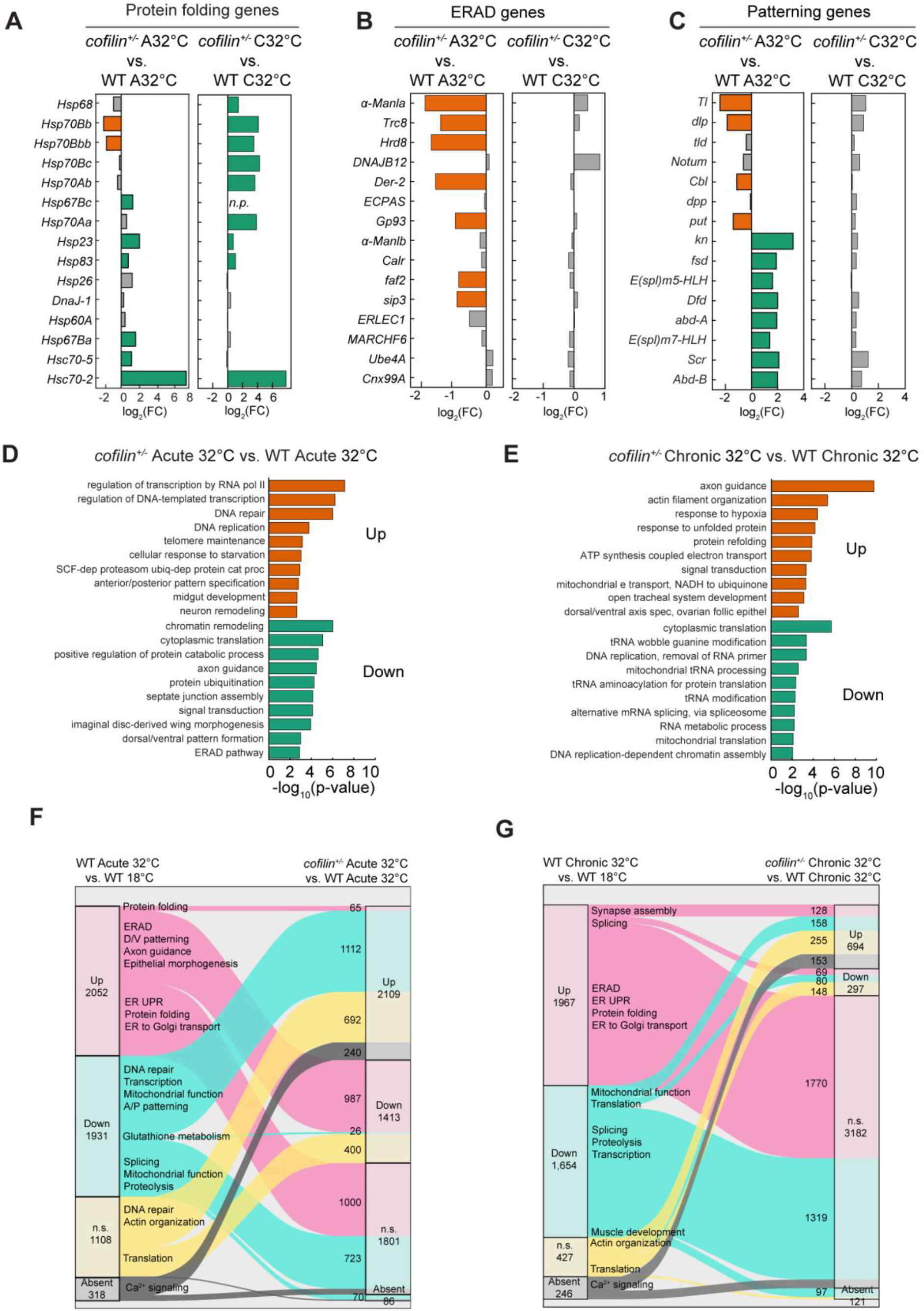
Heat-stressed *cofilin^+/-^* embryos show a distinct stress response compared to wild type. (A, B and C) Bar plots showing log_2_(FC) values for upregulated (orange) or downregulated (green) genes from highly significant ontologies related to: Protein folding (A), ERAD pathway (B) and Patterning (C) for the *cofilin^+/-^* acute 32°C versus (vs.) WT acute 32°C comparison (left) or the *cofilin^+/-^*chronic 32°C vs. WT chronic 32°C comparison (right). For Patterning, genes in the top 7 bars or bottom 8 bars are associated with dorsal/ventral or anterior/posterior patterning, respectively. Genes with no significant change in expression shown in gray. Abbreviation in (A) is: not present (n.p.). (D,E) Bar plots showing -log_10_(p-value) for selected highly significant GO Biological Processes for upregulated (orange) or downregulated (green) gene sets for the *cofilin^+/-^*acute 32°C versus vs. WT acute 32°C comparison (D) or the *cofilin^+/-^*chronic 32°C vs. WT chronic 32°C comparison (E). Abbreviated processes are: SCF-dependent proteasomal ubiquitin-dependent protein catabolic process; mitochondrial electron transport, NADH to ubiquinone; dorsal/ventral axis specification, ovarian follicular epithelium. (F, G) Alluvial plots showing change in direction of differential gene expression for genes in the WT acute 32°C vs. WT 18°C non-stressed comparison (left) relative to the *cofilin^+/-^* acute 32°C vs. WT acute 32°C comparison (right, F), or the WT chronic 32°C vs. WT 18°C non-stressed comparison (left) relative to the *cofilin^+/-^*chronic 32°C vs. WT chronic 32°C comparison (right, G). Selected significant ontologies shown for genes represented in corresponding ribbons. Number of genes up- (Up) or down-regulated (Down), not significant (n.s.) or Absent shown in outermost columns under the indicated comparisons. Number of genes per ribbon shown at right in middle column. Number of genes for very narrow ribbons not provided. Abbreviations are: dorsal/ventral (D/V); anterior/posterior (A/P); unfolded protein response (UPR). *Differential gene expression data in based on n = 6 biological replicates for 18°C WT and acute 32°C WT or n = 5 for chronic 32°C WT; and n = 6 for all conditions for cofilin^+/-^. Significance value cut-off for differentially expressed genes was FDR < 0.05; and significant ontologies p < 0.01*.

As a second approach to characterize the *cofilin*^+/-^ response with respect to wild type, we performed gene-to-gene comparisons. We asked the question: For either the acute or chronic heat stress exposure, if a gene is upregulated (or downregulated) in the wild-type stress versus non-stress comparison, what happens to the expression of the same gene in the *cofilin*^+/-^ versus wild-type comparison? This analysis again showed similarities between the two genotypes. For example, 1000 genes that were upregulated in wild-type embryos upon acute heat stress exposure did not change between *cofilin*^+/-^ and wild-type embryos under this condition, suggesting common responses (Fig 3F; S4 Table). These genes were enriched for processes related to protein folding and the ER unfolded protein response. For chronic stress, 1770 genes followed this pattern of behavior and were represented by some of the same biological processes plus ERAD (Fig 3G; S4 Table). Thus, evaluation at the gene-to-gene level shows that stress response in *cofilin*^+/-^ embryos is neither muted nor completely distinct from wild-type embryos and appears consistent with a reaction to reduced proteostasis.

However, we also noticed divergent regulatory directions for many genes, particularly under acute heat stress (Figs 3F and 3G; S3 Table). Some genes that were upregulated in wild-type embryos following acute heat stress were downregulated in the *cofilin*^+/-^ versus wild-type comparison (987 genes; Fig 3F). These genes were associated with ERAD, epithelial and neuronal tissue morphogenesis, and dorsal/ventral patterning among other biological processes (Fig 3F; S4 Table). Also, for the acute stress condition, an even larger set of genes that were downregulated in wild-type embryos were identified as upregulated in the *cofilin*^+/-^ to wild-type comparison (1112 genes; Fig 3F). These genes were involved in anterior/posterior patterning, transcription, DNA repair and mitochondrial function (Fig 3F; S4 Table). For both acute and chronic heat stress, genes associated with the actin cytoskeleton were not differentially regulated in wild-type embryos but were upregulated in *cofilin*^+/-^ embryos (Figs 3F and 3G; S4 Table). Finally, genes associated with cytoplasmic translation, which did not change in heat-stressed wild-type embryos, were downregulated in *cofilin*^+/-^ embryos for both heat stress conditions (Figs 3F and 3G; S4 Table).

Overall, a picture emerges that *cofilin*^+/-^ embryos respond to heat stress but with some modifications compared to the wild-type response. Signatures of pathways that manage reduced proteostasis are seen for both genotypes but with notable differences in induction of ERAD and changes in translation. At the same time, some heat compromised processes such as transcription seemed better maintained in *cofilin*^+/-^ versus wild-type embryos upon heat stress. We currently do not know which, if any, single divergent transcriptional response in *cofilin*^+/-^ embryos could account for their increased survival. Nonetheless, *cofilin*^+/-^ embryos display a transcriptional response to stress, while also moderating actin phenotypes associated with ASR; and together these actions correlate with enhanced survival.

### The transcriptome of *cofilin* knockdown embryos indicates stress prior to heat exposure

If *cofilin*^+/-^ embryos show indicators of stress response comparable to or exceeding wild type, then why was this not evident from the within-genotype comparison between non-stressed and heat-stressed *cofilin*^+/-^ embryos? The simplest explanation we could envision was that *cofilin*^+/-^ embryos already express a transcriptome indicative of stress response before encountering any elevated temperature condition.

To test this possibility, we compared our RNA-seq data for *cofilin*^+/-^ and wild-type embryos at the non-stressed 18°C condition. We found that 592 genes were significantly upregulated and 314 downregulated in the *cofilin*^+/-^ embryos (FDR < 0.05; Fig 4A; S3 Table). To confirm that changes in gene expression in non-stressed *cofilin*^+/-^ were indeed due to *cofilin knockdown*, we checked for reproducibility using a different mutant allele of *cofilin* (*tsr^N121^*). Based on qPCR results, maternal heterozygosity for *tsr^N121^* led to roughly the same knockdown in *cofilin* transcript level in embryos as did maternal heterozygosity for *tsr^1^* (S2 Table). We found that for four genes that were highly upregulated following knockdown using the *tsr^1^* allele, three were similarly upregulated by knockdown with *tsr^N121^* (S2 Table), supporting that the changes in gene expression in non-stressed *cofilin*^+/-^ embryos result from the reduced dosage of Cofilin.

**Fig 4.**
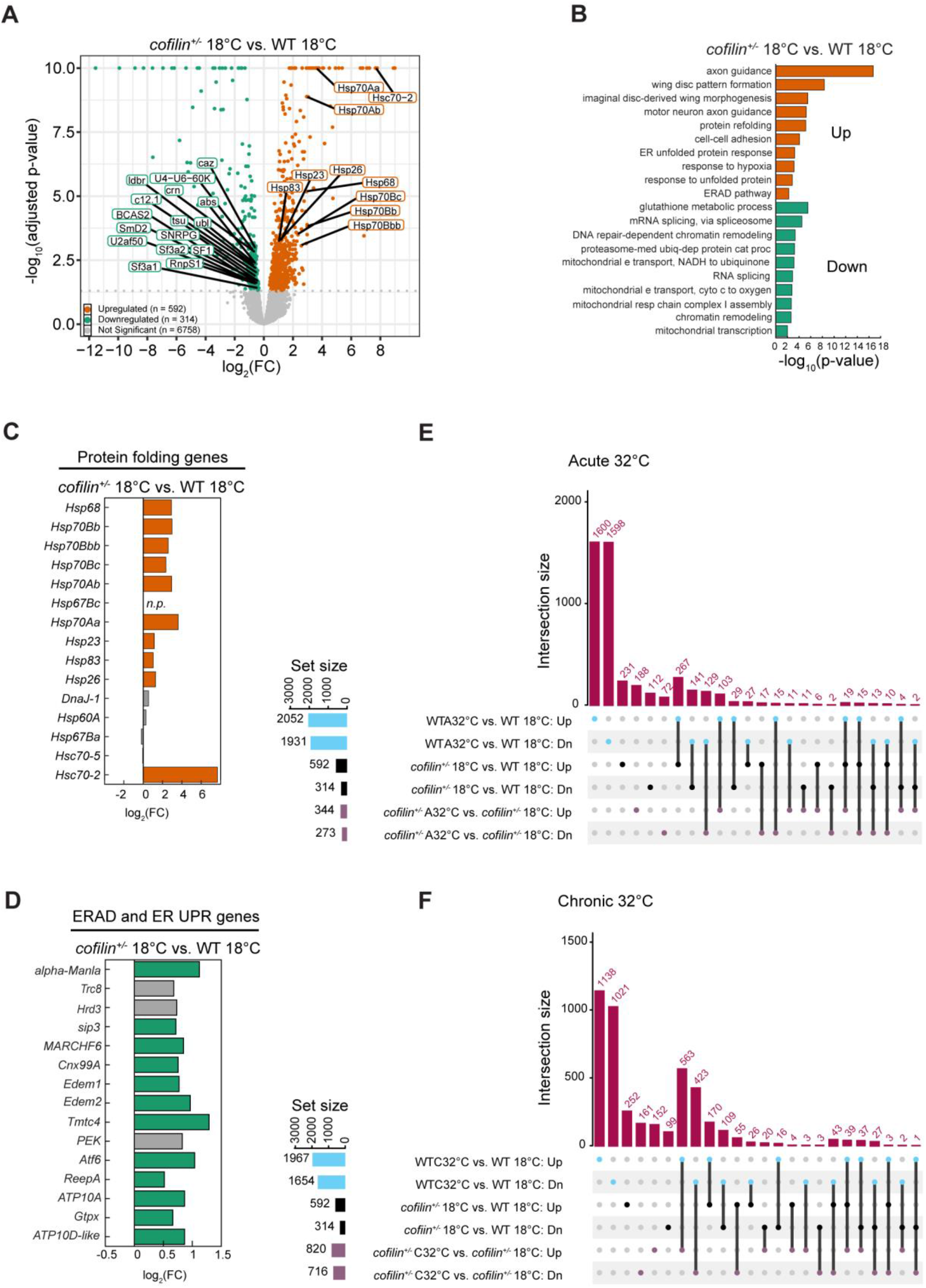
The *cofilin^+/-^* embryos show signs of stress response absent any heat stress exposure. (A) Volcano plot showing -log_10_(adjusted p-value) versus log_2_(FC) for upregulated (orange) and downregulated (green) genes in 18°C non-stressed *cofilin^+/-^* embryos versus (vs.) 18°C non-stressed WT embryos. Number of genes up- or down-regulated, or not significant shown in key. Select HSPs and splicing genes labeled for the up- or down-regulated genes, respectively. Genes associated with the processes of protein folding or splicing chosen due to their common direction of expression observed for most comparisons performed. (B) Bar plot showing -log_10_(p-value) for selected highly significant GO Biological Processes for upregulated (orange) or downregulated (green) gene sets for 18°C non-stressed *cofilin^+/-^* embryos versus (vs.) 18°C non-stressed wild-type (WT) embryos. Abbreviated processes are: proteasome-mediated ubiquitin-dependent protein catabolic process; mitochondrial electron transport, NADH to ubiquinone; mitochondrial electron transport, cytochrome c to oxygen; mitochondrial respiratory chain complex I assembly. (C, D) Bar plots showing log_2_(FC) values for upregulated (orange) or downregulated (green) genes from highly significant ontologies related to: Protein folding (C) and combined ERAD pathway and ER unfolded protein response (D) for 18°C non-stressed *cofilin^+/-^* embryos vs. 18°C non-stressed WT embryos. Genes with no significant change in expression shown in gray. Abbreviations are: not present (n.p.), unfolded protein response (UPR). (E, F) UpSet plots showing intersection between up- or down-regulated gene sets from pairwise comparisons for acute (E) or chronic (F) 32°C heat-stressed embryos. Comparisons as indicated. Numbers in black at left indicate how many differentially expressed genes are included per set. Vertical black lines connecting dots indicate gene sets that intersect, and magenta bars and numbers indicate how many differentially expressed genes are shared. Abbreviations are: Down (Dn); Acute (A); Chronic (C). *Differential gene expression data in (A-F) based on n = 6 biological replicates for 18°C WT and 18°C cofilin^+/-^. Significance value cut-off for differentially expressed genes was FDR < 0.05; and significant ontologies p < 0.01*.

Interestingly, the upregulated genes in non-stressed *cofilin*^+/-^ embryos were involved in many processes related to neuronal and epithelial tissue building, as we saw previously for the upregulated genes in wild-type embryos exposed to either acute or chronic heat stress (compare Figs 2B and 2C to Fig 4B; S4 Table). Furthermore, several HSP genes, including all the Hsp70s, Hsps 68, 23, 26 and 83, and Hsc70-2 were upregulated (Fig 4C; S3 Table). Processes related to ERAD and ER unfolded protein response were also identified for the upregulated genes in non-stressed *cofilin*^+/-^ embryos (Figs 4B and 4D; S4 Table). Approximately 45% of the upregulated genes in non-stressed *cofilin*^+/-^ embryos were shared with upregulated genes in wild-type embryos exposed to acute stress (267/592 genes; for chronic stress, 29% or 170/592 genes; Figs 4E and 4F). Thus, cofilin knockdown embryos sense some form of stress, perhaps related to loss of proteostasis, even in the absence of heat exposure.

The downregulated genes for non-stressed *cofilin*^+/-^ embryos were involved in splicing, DNA repair, and mitochondrial activity, again consistent with the downregulated genes identified for wild-type embryos in response to heat stress (compare Figs 2B and 2C to Fig 4B; S4 Table). Once more, we found that 45% of the downregulated genes in non-stressed *cofilin*^+/-^ embryos were shared with downregulated genes in wild-type embryos exposed to acute stress (141/314 genes; for chronic stress, 35% or 109/314 genes; Figs 4E and 4F). Together, our results show that *cofilin* knockdown alone leads to transcriptomic remodeling prior to any heat-stress challenge. These embryos are already experiencing moderate stress due to the reduced genetic dosage of *cofilin*. The alternate *cofilin*^+/-^ baseline resembles a reduced version of heat stress response in wild-type embryos, with strong indications of reduced proteostasis in these embryos. We suggest that this alternate baseline in non-stressed *cofilin*^+/-^ embryos represents an adapted state that promotes increased survival once the embryos do encounter heat stress.

## Discussion

We initiated our study with a focus on Cofilin’s role in driving cytoplasmic actin destabilization and nuclear rod formation as part of the ASR in heat-stressed embryos. We found that reducing Cofilin dosage attenuates these actin disruptions and improves embryo survival under both acute and chronic stress conditions. However, our data do not support the view that improved survival results only from the suppression of the observed actin phenotypes. Instead, we found that reducing Cofilin levels introduces moderate physiological stress on its own, such that *cofilin*^+/-^ embryos already display signs of heat shock and the ER unfolded protein response, even in the absence of heat stress. Once *cofilin*^+/-^ embryos are exposed to heat stress, their response appears muted or distinct compared to wild-type embryos, but this is largely because they already express stress related genes. Coincidentally, in this adapted state, *cofilin*^+/-^ embryos show increased heat tolerance.

Coincident with the ASR, our RNA-seq approach revealed upregulation of genes associated with heat shock response, the ER unfolded protein response and the ERAD pathway. Early *Drosophila* embryos are already known to mount some limited aspects of heat shock response – specifically, HSP expression – although a full response is not possible until later in development (60, 62, 63, 100–102). Our results show that several HSPs and related chaperones are upregulated even at 32°C, well below the 37-42°C range, which is commonly used in heat shock experiments (60–63). More unexpectedly, we found induction of genes related to ER stress response and ERAD in heat-stressed embryos. We are not aware of previous reports of induction of ER stress response pathways in fly embryos. However, ER stress response is readily activated in mammalian pre-implantation embryos, where it compromises embryo survival and has important consequences for both agricultural and clinical applications (103–106). Together, the concurrence of heat shock response and ER stress response implies that heat-stressed fly embryos sense and mount a transcriptional response to perturbed proteostasis, suggesting that this tractable model system may be well-suited for further study of the interplay between protein homeostasis and development.

We found that *cofilin*^+/-^ embryos show a small but statistically significant increase in survival upon heat stress. This increased resilience of *cofilin*^+/-^ embryos may represent the biological phenomenon of hormesis. Hormesis is a protective state, whereby a mild or moderate initial stress leads to improved outcomes when a biological system encounters subsequent or more severe stress (107). Hormesis is well documented in both embryos and adult organisms and promotes cell survival as well as organismal longevity (108–114). Inducers of hormesis may include genotypic, mitochondrial, oxidative, heat or osmotic stressors. In one simple mechanistic scenario, the priming stress pre-loads cells with HSPs that then aid future buffering responses (63, 102, 111, 115). Alternatively, priming can be accomplished through complex mechanisms such as a mild stressor driving changes in gene expression programs that provide long-lasting adaptation (e.g., mild mitochondrial dysfunction leading to nuclear-dependent remodeling of metabolism) (110, 116–118). In *cofilin*^+/-^ embryos, the former mechanism is plausible because our RNA-seq data show evidence of the expression of stress response proteins in the non-stressed embryos. In this case, maternal preloading of transcripts and/or proteins could be particularly beneficial in chronically heat-stressed early embryos, as these embryos may have limited capacity to respond to stress until the zygotic genome is fully activated (63, 101, 111, 119). Our gene expression analysis also suggests that this adaptation makes *cofilin*^+/-^ embryos better able to maintain core cellular processes such as transcription, DNA repair and mitochondrial activity when they later encounter heat stress.

It is not clear why the *cofilin*^+/-^ genotype induces physiological stress in embryos in the absence of an external stressor. We did note that *cofilin*^+/-^ embryos show slightly reduced survival at the non-stress temperature, as well as slower F-actin turnover. This suggests that reduced Cofilin dosage is not sufficient to support normal F-actin dynamics, and so any of the numerous actin-dependent events that Cofilin contributes to could be compromised. Thus, *cofilin* knockdown could somehow indirectly precipitate mild cellular stress.

One striking observation is that the stress response triggered by *cofilin* heterozygosity indicates a loss of proteostasis, as indicated by upregulation of HSPs and ER unfolded protein response pathways. Possible links between the actin cytoskeleton and proteostasis are numerous since actin could conceivably impinge on almost any aspect of mRNA expression, mRNA metabolism, or protein production, degradation or distribution (120–122). For example, nuclear actin regulates nucleolar structure, RNA Pol I transcription, ribosomal RNA synthesis and, consequently, global translation (123, 124). Also, specific F-actin structures have been described as functionally important translation platforms in organisms spanning from yeast to mammals (e.g. see (125–130)). Translation factors, including both initiation and elongation factors, interact with F-actin (121). For eEF1A, its mutually exclusive binding with either F-actin or amino acylated tRNAs provides a bridge between actin and translational regulation (121, 131–134). Regarding protein degradation, proteasome activity is associated with F-actin remodeling (129); and clearance of dysfunctional proteins by autophagy inside the cell and by endocytic retrieval at the cell surface requires F-actin (135, 136). Cofilin could then interact with proteostasis at any of these steps via its control over F-actin dynamics (137).

Thus, our findings reveal yet another possible intersection between Cofilin biology, inducible stress response and survival: Here, Cofilin appears to modulate protein homeostasis, which is central to cellular stress response. Prior studies have repeatedly shown that reducing Cofilin levels or activity promotes the survival of stressed mammalian cells to the extent that Cofilin reduction is being pursued as a therapy for neurodegeneration and ischemic tissue injury (18, 46–48, 138–142). In those contexts, the survival benefit of Cofilin reduction has been assigned to multiple mechanisms. For example, reduced Cofilin increases neuronal viability by limiting the assembly of actin rods that physically impede transport along axons and dendrites (34, 35, 44–46). Alternatively, reducing Cofilin can increase survival in chronic stress conditions by blocking programmed cell death pathways that are activated by highly oxidized Cofilin (5, 6, 141, 143). Now, from fly embryos, we add that low-grade stress resulting from *cofilin* knockdown and an associated reduction in proteostasis may also enhance subsequent stress response and survival outcomes through adaptive mechanisms, such as hormesis.

In our study of embryos, we found no immediate support for the protective value of the ASR. It has been argued that when stress is transient, actin rods can be beneficial because they prevent actin, an ATPase, from consuming unnecessary energy as part of the polymerization cycle (43, 49). Counter to this, when we inhibited rod formation in our acute heat stress condition, the embryos showed improved stress tolerance. However, considering our observations regarding baseline stress in the *cofilin*^+/-^ embryos, and the potential for stress adaptation, we are reluctant to interpret our results from the perspective of rods alone. Instead, our systems-wide survey emphasizes that a complex relationship exists between Cofilin, multiple stress response pathways, and perhaps proteostasis. Given these interactions, a major question that emerges is whether there are primary and direct Cofilin activities that benefit cells as they try to cope with short- or long-term physiological stress. In one simple model, Cofilin may somehow normally promote protein quality control, and under stress Cofilin is activated to facilitate the clearance of unfolded proteins.

In this scenario, ASR and rod formation may be more a by-product of protective stress response rather than its goal.

## Materials and methods

### Resource availability

Requests for information or resources should be directed to the corresponding author, Anna Marie Sokac.

### Materials availability

All materials used in this study are commercially available from the sources listed in the key resources table or available upon request from the lead contact. Fly stocks are available from Bloomington *Drosophila* Stock Center (Indiana University, Bloomington, IN, USA).

### Data and code availability

- Raw RNA-seq data will be available at the NIH NCBI (https://www.ncbi.nlm.nih.gov/bioproject/) under BioProject Number: PRJNA1368972. *Data will be released upon publication*.
- Raw mass spectrometry data is available at the PRIDE database (https://www.ebi.ac.uk/pride/) under Project Accession: PXD058671; Token: dDxylt5Mqa8L.
- Raw and analyzed imaging data will be available at the Illinois Data Bank (https://databank.illinois.edu/datasets) under dataset ID: 2698771. *Data will be released upon publication*.
- Any additional information required to reanalyze the data in this study is available from the lead contact upon request.

### Experimental model and subject details

*Drosophila melanogaster* stocks were housed on Fly Food R (Lab-Express, Ann Arbor, MI, USA). Analyses in this study focus on cellularization, corresponding to Bownes Stage 5 (78) unless otherwise noted in the Method Details. We used FlyBase to retrieve gene sequences and information regarding *Drosophila* reagents (144).

### Fly stocks and genetics

For imaging and mass spec experiments, Oregon R (Ore-R) was used as the wild-type stock. For all other experiments, the *cofilin^+/-^*embryos were prepared by crossing 3–5-day old virgin female flies from *tsr^1^/CyO* or *tsr^N121^/CyO* (BDSC#9107 or BDSC#9109, respectively; Bloomington Drosophila Stock Center), with 3–5-day old males from Ore-R at 25°C. F1, 3–5-day old, *tsr^1^/+* (or *tsr^N121^/+*) virgin females were then crossed with 3–5-day old sibling males (for imaging, RNA-seq and qPCR) or Ore-R males (for survival assays) to generate F2 embryos for experiments. The outcross “wild-type” control embryos were prepared according to the same strategy as the *cofilin^+/-^* embryos, except that the initial parental cross was between *white^1118^* females and Ore-R males.

### Method details

#### Embryo heat stress exposure

Embryos were harvested from apple juice agar plates affixed to collection cups as previously described in Biel 2020 (50). To allow flies to acclimate to the starting temperature of either 18°C (for non-stressed and acute heat stress conditions) or 32°C (for the chronic heat stress condition), collection cups were incubated at 18°C for 48 hours or 32°C for 12 hours. The acclimation period at 32°C was shorter because extended incubation caused dramatically reduced fertility and increased fly death, such that these cups were always discarded after 24 hours. After the acclimation period, embryo collections proceeded according to the schematic in Fig 1A. Specifically, for the 18°C non-stressed condition, flies were allowed to lay eggs for 2 hours, then plates were incubated at 18°C for an additional 3.5 hours. For the acute 32°C heat stress condition, flies were allowed to lay eggs at 18°C for 2 hours, then plates were incubated at 18°C for an additional 20 minutes, after which they were heat stressed at 32°C for 1.5 hours. Embryo temperature reaches 32°C ∼10 minutes after moving plates between incubators. For the chronic 32°C heat stress condition, flies were allowed to lay eggs at 32°C for 1 hour, then plates were incubated at 32°C for an additional 1.5 hours. Embryo cups at 32°C were discarded after 24 hours. Embryos were batch-collected or hand-selected as indicated per experiment.

#### Survival assays

For the larval hatching assay, embryos with intact chorion were loosened from apple juice agar plates with distilled water and a paintbrush, then batch-collected in a cell strainer and washed thoroughly with distilled water. Embryos were transferred to a fresh agar plate with a paintbrush and hand-selected at cellularization stage under Halocarbon 27 oil (MilliporeSigma, Burlington, MA, USA). Selected embryos were aligned in rows on fresh agar plates, placed in a plastic box with damp paper towels, and kept at room temperature (22°C) with controlled humidity of 40% for 48 hours before empty embryo eggshells and dead embryos were counted.

#### Immunostaining

Embryos were batch-collected from apple juice plates by dechorionation with 3% bleach and fixed in 4% paraformaldehyde in 1X PBS according to previously published methods (51). Fixed embryos were washed in methanol and stored at -20°C for a minimum of 24 hrs.

For immunostaining, plasma membrane furrows were detected with anti-Peanut primary antibody (1:200) (DSHB, Iowa City, IA, USA) and goat anti-mouse Alexa Fluor 488 secondary antibody (1:500) (Thermo Fisher Scientific, Waltham, MA). Nuclei were visualized with Hoechst 33342 dye (1.0 μg mL^-1^) (Thermo Fisher Scientific).

Prior to imaging, embryos were mounted on slides in Aqua Poly/Mount (Polysciences, Inc., Warrington, PA, USA) and covered using a 1.5 mm coverslip (Corning, Corning, NY, USA).

#### Rhodamine G-actin injections

Embryos were batch-collected from apple juice plates by dechorionation with 3% bleach at 18°C or 32°C and prepped for injection as in Biel, et al., 2020, at least 30 minutes prior to cellularization. All reagents and equipment were pre-chilled or pre-warmed to maintain temperatures to the greatest extent possible. Lyophilized rhodamine-conjugated non-muscle human platelet G-actin (G-actin^Red^, Cytoskeleton Inc., Denver, CO, USA) was prepared by adding 1 μL of nuclease free water and 1 μL of freshly prepared G-buffer (5 mM Tris-HCl, 0.2 mM CaCl_2_, 1 mM DTT, 0.2 mM ATP, pH 8.0) to 10 mg of G-actin^Red^. A glass capillary needle was loaded with 1.5 μL of the prepared G-actin^Red^ and ∼50 pL injected per embryo (32, 50). After injection, embryos were returned to their respective temperature conditions to incubate in humidified chambers until imaging at cellularization.

#### Image acquisition and presentation

Images were collected on a Zeiss LSM 880 Airyscan confocal microscope using a Plan-Apochromat 63x oil immersion objective with 1.4 numerical aperture for fixed imaging or C-Apochromat 40x water immersion objective with 1.2 numerical aperture for live imaging (Carl Zeiss, Inc., Oberkochen, Germany).

For furrows and multinucleation, images were collected as surface views from fixed wild-type or *cofilin^+/-^*embryos at focal planes ∼2-10 μm beneath the embryo surface. Images were acquired as 1024×1024 pixels/frame with 7.5879 pixels/micron for quantifying multinucleation or 15.1759 pixels/micron for presentation. For each embryo a corresponding cross-sectional image was collected to allow measurement of the plasma membrane furrow length, which serves as a proxy for time during cellularization (70). Images for analysis and presentation were limited to the same time window within cellularization. For presentation, color channels were adjusted separately, noise removed using the Despeckle algorithm in ImageJ/Fiji (NIH; Bethesda, MD, USA), and a Gaussian blur filter of 2 was applied in Adobe Photoshop CC (Adobe, San Jose, CA, USA).

For FRAP of cytoplasmic furrow tips, living WT or *cofilin^+/-^* embryos were imaged after G-actin^Red^ injection at 18 ± 2°C or 32 ± 2°C in a thermal incubator. For the 32°C condition, the objective was also heated to 32 ± 2°C using an objective heater consisting of a thermal collar and temperature controller (Okolab, Sewickley, PA, USA). G-actin^Red^ at furrow tips was bleached to approximately 60-70% of initial fluorescence intensity using the 561 nm laser at 100% power in cross-section views at the embryo equator. The size of the bleached box was 1.25×1.25 μm. Fluorescence recovery was tracked for 90 seconds at 1 second intervals. Images were acquired as 1024×1024 pixels/frame with 4.8177 pixels/micron, but the acquisition region was limited to 100×100 pixels to allow fast acquisition. G-actin^Red^ signal from an unbleached neighboring furrow tip was used to ascertain background bleaching due to imaging. Bleached furrows ingressed at the same rate as unbleached furrows arguing that bleaching did not cause phototoxicity (32, 70). For presentation, images were cropped, contrast adjusted, and a Gaussian blur of 0.5 was applied in Adobe Photoshop CC.

For nuclear actin rods, living wild-type or *cofilin^+/-^*embryos were imaged at 18 ± 2°C or 32 ± 2°C in a thermal incubator as described for furrow tip FRAP. For the 32°C condition, the objective was also heated to 32 ± 2°C using an objective heater. Z-series surface views were collected at ∼2-10 μm beneath the embryo’s surface, within the top two-thirds of the nuclei. Images were acquired as 1024×1024 pixels/frame with 3.6133 pixels/micron for counting rods or 9.6355 pixels/micron for quantifying rod morphology. For presentation, actin rod images were adjusted for brightness, cropped in ImageJ/Fiji and a Gaussian blur filter of 0.75 was applied in Adobe Photoshop CC.

For FRAP of actin rods, living WT or *cofilin^+/-^* embryos were imaged after G-actin^Red^ injection as described for furrow tip FRAP with the following exceptions: Actin rods were imaged in surface views at ∼2-10 μm beneath the embryo’s surface. The size of the bleached box was 0.2×0.2 μm. Fluorescence recovery was tracked for 60 seconds at 1-2 second intervals. Images were acquired as 1024×1024 pixels/frame with 9.6355 pixels/micron, but the acquisition region was limited to 100×100 pixels or less for faster acquisition. Cytoplasmic G-actin^Red^ signal from an unbleached neighboring region was used to ascertain background bleaching due to imaging. For presentation, images were cropped, contrast adjusted, and a Gaussian blur of 0.5 was applied in Adobe Photoshop CC.

#### RNA sequencing

Embryos were batch-collected from apple juice plates as for the survival assays, using water, at 18°C or 32°C. Reagents and equipment were pre-chilled or pre-warmed to maintain temperatures to the greatest extent possible. Embryos were transferred to a fresh agar plate with a paintbrush and cellularization stage embryos selected under Halocarbon 27 oil (MilliporeSigma). Selected embryos were snap-frozen using liquid nitrogen and stored at -80°C.

RNA was extracted from six biological replicates of embryos collected from outcrossed wild-type and *cofilin^+/-^* mothers per temperature condition. In addition, two more biological replicates of wild-type (Ore-R) fly embryos were included as batch controls during collection of both outcrossed wild-type and *cofilin^+/-^* embryos to address possible batch associated variations. RNA was extracted using the RNeasy Plus Micro Kit (Qiagen, Germantown, PA, USA). RNA quality was confirmed by measuring the RNA Quality Number & 28S/18S rRNA ratio using AATI Fragment Analyzer (Advanced Analytics, Ames, IA, USA).

Construction of the stranded RNA-seq libraries and sequencing were performed at the Roy J. Carver Biotechnology Center (University of Illinois at Urbana-Champaign). Purified DNase-treated total RNAs were run on a Fragment Analyzer (Agilent, Santa Clara, CA, USA) to evaluate RNA integrity. The total RNAs were converted into individually barcoded polyadenylated mRNAseq libraries using the KAPA HyperPrep mRNA Kit (Roche, Indianapolis, IN, USA) following the manufacturer’s protocol. The final libraries were quantified with Qubit (Thermo Fisher Scientific), and the average cDNA fragment sizes were determined on a Fragment Analyzer. The libraries were pooled and further quantified by qPCR on a CFX Connect Real-Time qPCR system (Bio-Rad, Hercules, CA, USA) to ensure maximization of cluster numbers in the flowcell.

The pooled barcoded RNA-seq libraries were sequenced using a NovaSeq X Plus (for outcrossed wild-type embryos) or a NovaSeq 6000 (for *cofilin^+/-^*embryos) (Illumina, San Diego, CA, USA). Two independent inbred Ore-R control libraries were included in each sequencing run to assess potential sequencing-run-associated variation between the NovaSeq instruments. The libraries were sequenced from both ends of the fragments for a total of 150 bp per end. The fastq read files were generated and demultiplexed using the Illumina demultiplexing software (Illumina).

#### qPCR

Wild-type embryos were collected for three biological replicates per condition, and RNA was extracted as for RNA-seq. Total RNA samples of 300ng were reverse transcribed using oligo (dT) primers and the iScript Select cDNA Synthesis Kit (Bio-Rad). cDNA was diluted 1:5 and real-time qPCR performed on the Applied Biosystems StepOnePlus Real-Time PCR system using TaqMan Fast Advanced Master Mix and TaqMan Gene Expression Assays (Applied Biosystems / Thermo Fisher Scientific). Predesigned TaqMan Gene Expression Assays (Thermo Fisher Scientific) were used for *Hsp23* (Assay ID: Dm01822473_s1), *Hsp26* (Assay ID: Dm01822452_s1), *Hsp68* (Assay ID: Dm02151262_s1), *Hsc70-4* (Assay ID: Dm02153823_s1), *Hsp83* (Assay ID: Dm02362342_s1), and *DnaJ-1* (Assay ID: Dm01832926_s1), *CG15894* (Assay ID: Dm01833202_g1), *CG6180* (Assay ID: Dm01790866_s1), *CG11034* (Assay ID: Dm01804861_g1), *CG6282* (Assay ID: Dm01841146_m1), and tsr (Assay ID: Dm01842418_g1). A custom TaqMan Gene Expression Assay (Thermo Fisher Scientific) was used for *Hsp70*. The gene *roadblock* (*robl*; Thermo Fisher Scientific, Catalog# 4351372, Assay ID: Dm01821522_g1) served as a reference control to normalize Cτ values per HSP gene. The *robl* gene was selected due to its stable expression across experimental groups in the RNA-seq experiments.

#### Western blotting

Wild-type and *cofilin^+/-^* embryos were collected as for RNA-seq. Three biological replicates of 200 embryos per condition were homogenized by hand on ice in lysis buffer (150 μL 0.05 M Tris-HCl pH 8.0, 0.15 M KCl, 0.05 M EDTA, 0.5% NP-40, 1X protease inhibitor cocktail (Thermo Fisher Scientific)) in low-protein retention microcentrifuge tubes (Thermo Fisher Scientific) using sterile plastic pestles. Embryo debris was pelleted, and soluble lysate collected. Protein concentration was determined by BCA assay (Thermo Fisher Scientific) and lysates diluted with lysis buffer to obtain the same concentration between samples. Lysates were boiled with 1x SDS-PAGE reducing buffer (62.5 mM Tris-HCl pH 6.8, 2% SDS, 10% glycerol, 41.6 mM DTT, 0.01% bromophenol blue) and run at a concentration of 5 μg per lane through 12% hand cast bis-acrylamide separating gels. Protein was transferred to 0.2 μm nitrocellulose (Bio-Rad) and probed with 1:200 mouse anti-β-actin (Santa Cruz Biotechnology, Dallas, TX, USA) and 1:5000 rabbit anti-*Dm*-Cofilin (32), followed by goat anti-mouse or goat anti-rabbit HRP secondary antibodies (Jackson ImmunoResearch Labs, West Grove, PA, USA).

#### Mass spectrometry

Embryo collection and protein lysate preparation followed the same methodology as for Western Blotting with the following exceptions: Approximately 62 μg of protein was loaded per gel lane and run through pre-made 4-20% bis-polyacrylamide SDS-PAGE separating gels (Bio-Rad) beside pre-stained molecular weight standards (MilliporeSigma). Lysates were loaded with empty lanes separating them to prevent contamination between samples. Gels were fixed and stained with QC Colloidal Coomassie G-250 stain according to the manufacturer’s Quick Stain protocol (Bio-Rad) and gel regions between the 17-35 kDa and 50-100 kDa molecular weight markers were cut out per lane with a clean razor blade. Gel pieces were stored and shipped in low-protein retention tubes at 4°C.

Gel pieces were washed with 50 mM ammonium Bicarbonate (NH_4_HCO_3_; Acros Organics) in 50% acetonitrile (ACN; Fisher Scientific, Hampton, NH, USA) and dehydrated with 100% ACN. The gel pieces were reduced with 10 mM Tris (2-carboxyethyl) phosphine (TECP; MilliporeSigma) at 55°C for 30 minutes, followed by alkylation with 10 mM 2-chloroacetamide (CAA; MilliporeSigma) for 45 minutes at room temperature in the dark. Then, gel pieces were dehydrated in ACN and dried prior to overnight digestion with a 20 ng/µl trypsin (Thermo Fisher Scientific) solution prepared in 50 mM NH_4_HCO_3_. Digestion was halted with 5% formic acid (FA; Fisher Scientific) and the solution transferred to clean tubes. Peptides were extracted from the gel pieces by incubation with 0.1% FA in 50% ACN for 15 minutes, and the extracted solution was combined with the previous digestion solution. Digested peptides were desalted using ZipTip pipette tips (MilliporeSigma) according to the manufacturer’s protocol. After desalting, peptides were dried and reconstituted in water containing 0.2% FA and 2% ACN for subsequent LC-MS/MS analysis.

Mass spectrometry was performed with a Vanquish Neo UHPLC system (Thermo Fisher Scientific) coupled to an Orbitrap Eclipse Tribrid mass spectrometer (Thermo Fisher Scientific). Peptides were separated on an Aurora UHPLC Column (25 cm × 75 μm, 1.7 μm C18, AUR3-25075C18-TS; Ion Opticks, Collingwood, VIC, Australia) with a flow rate of 0.35 μL/min for a total duration of 43 min and ionized at 1.6 kV in the positive ion mode. The gradient was composed of 6% solvent B (3 min), 6-25% B (20 min), 25-40% B (7 min), and 40–98% B (13 min); solvent A: 0.1% (v/v) FA; solvent B: 0.1% (v/v) FA/80% (v/v) ACN. MS1 scans were acquired at the resolution of 120,000 from 350 to 2,000 m/z, AGC target 1e6, and maximum injection time 50 milliseconds. MS2 scans were acquired in the ion trap using fast scan rate on precursors with 2-7 charge states and quadrupole isolation mode (isolation window: 1.2 m/z) with higher-energy collisional dissociation (HCD, 30%) activation type. Dynamic exclusion was set to 30 seconds. The temperature of ion transfer tube was 300°C and the S-lens RF level was set to 30.

### Quantifications and statistical analysis

Student’s t-tests were performed using custom code from MATLAB’s Statistics and Machine Learning Toolbox (Mathworks, Natick, MA, USA). Chi-square contingency analysis was performed using GraphPad QuickCalcs (GraphPad, San Diego, CA, USA). Specific p-values and n-values can be found within the figures and their legends. For imaging data, comparisons with p < 0.05 were considered significant. For RNA-seq, genes following the significance cutoff clearance: FDR < 0.05 were considered differentially expressed (up- or down-regulated). For identifying important ontologies, GO Biological Processes following the significance cutoff clearance: p ≤ 0.01 were considered. All plots were generated using custom codes in MATLAB, except for the volcano and UpSet plots which were generated using R (Vienna, Austria) packages ‘ggplot2’ and ‘UpSetR’, respectively. Final plots and figures were assembled using Adobe Illustrator CC.

#### Actin rod quantification

To quantify the percent embryos displaying actin rods in Fig 1G, surface view images with ≥ 250 visible nuclei were scored manually, and the presence of rods was defined as bright G-actin^Red^ streaks visible within the nuclei, approximately 0.1-0.2 μm in diameter and present in at least two contiguous focal planes.

To quantify rods per nucleus in Fig 1H, rods were identified manually in ImageJ/Fiji from a 256×256 pixel region of the raw Z-stack (∼80-130 nuclei counted per embryo), scrolling through multiple focal planes to assure accurate counting. Rods and nuclei were then counted individually using the Selection and Measure tools, and the ratio of rods/nuclei calculated per embryo.

To quantify rod diameters in S1C Fig, five nuclei representing four quadrants and a central region of a 512×512 pixel cropped image from the raw Z-stack were chosen. The diameters of all rods in each nucleus were measured manually using the Line and Measure tools in ImageJ/Fiji. If one nucleus contained more than one rod, the mean diameter for that nucleus was calculated. The five means per embryo were plotted as individual data points in the final figure.

To quantify FRAP for rods in S1E Fig, mean fluorescence intensity from the 0.2×0.2 μm bleached region was manually measured for each frame using ImageJ/Fiji. Fluorescence intensity from a 0.2×0.2 μm unbleached cytoplasmic region was also measured for each frame to account for photobleaching effects. Half-time to fluorescence recovery (t_1/2_) was calculated in MATLAB using Method 1 described in Figard et al., 2019 (32). MATLAB was used to generate the recovery fit curves.

For all rod experiments, t_1/2_ recovery curve data shown is based on n ≥ 3 independent biological replicates.

#### Furrow tip F-actin FRAP analysis

To quantify actin furrow tip FRAP in Fig 1E, mean fluorescence intensity from the 1.25×1.25 μm bleached region was manually measured for each frame using ImageJ/Fiji. Fluorescence intensity from a 1.25×1.25 μm region in the unbleached cytoplasm neighboring the bleached furrow was also manually measured for each frame to account for photobleaching effects. Half-time to fluorescence recovery data (t_1/2_) in Fig 1E and percent mobile fraction data in S1B Fig were calculated in MATLAB using Method 1 described in Figard et al., 2019 (32).

For all furrow tip FRAP experiments, t_1/2_ data and corresponding percent mobile fraction data shown is based on n ≥ 3 independent biological replicates.

#### Multinucleation quantification

To quantify the percent embryos with multinucleation, raw z-stack images were scored manually for the presence of multinucleation (≥100 nuclei per image), scanning through multiple focal planes to assure accurate counting.

To quantify the ratio of multinucleate cells/total nuclei in Fig 1C, the number of multinucleated cells and total nuclei in the frame of each image were counted manually in ImageJ/Fiji using the Pencil and Multi-point selection tools. Segmented images were generated from the raw images manually using the Pen tool in Adobe Illustrator CC.

For all multinucleation experiments, data shown is based on n ≥ 3 independent biological replicates. Trends are consistently observed, but a chi-square contingency test for the binary multinucleation phenotype and a Student’s t-test for the multinucleation ratio do not show statistical significance. A complementary exact Cochran–Mantel–Haenszel (CMH) analysis of the multinucleation frequency for combined acute and chronic heat-stress conditions does show a statistically significant difference between genotypes (p = 0.0149). Nonetheless, we present the ratios as only descriptive in Fig 1C.

#### Survival assay statistical analysis

Embryo hatching assays were performed in 3 independent biological replicates per condition, with 200 embryos scored per replicate. The reported n=600 represents the total number of embryos pooled across all independent replicates. For each replicate, percent survival was calculated as the number of embryos that hatched as live larva divided by the total number of embryos scored. Statistical significance was assessed for the raw hatched and unhatched embryo counts across conditions using a chi-square contingency test with a two-tailed p-value.

#### RNA-seq analysis

Raw RNA-seq data was checked for read quality using FastQC (version 0.11.9; Babraham Bioinformatics, Cambridge, UK). Reads were aligned to the *Drosophila* genome from FlyBase (FASTA: dmel-all-chromosome-r6.50.fasta, GTF: dmel-all-r6.50.gtf) using STAR (version 2.7.11b) (145). Sorted bam files were then indexed using SAMtools index. Gene-level read counts were generated using HTSeq-count (version 2.0.2) in BAM mode (-f bam) with position-sorted alignments (-r pos) and reverse-strand specificity (-s reverse), consistent with the stranded KAPA mRNA HyperPrep library preparation protocol (146). Read normalization, variance estimation, and pairwise differential expression analysis was performed using the Bioconductor package EdgeR (version 3.40.2) (147). For each pairwise comparison, genes with counts per million (CPM) > 1 in at least two samples across the groups being compared were retained. Library-size normalization was performed using the trimmed mean of M-values (TMM) method. A no-intercept design matrix was constructed for each comparison, and dispersions were estimated using estimateDisp with robust estimation enabled. Gene-wise negative-binomial generalized linear models were fitted using glmFit, and pairwise contrasts were tested using likelihood-ratio tests with glmLRT. Genes with FDR < 0.05 were considered differentially expressed. Gene set enrichment analysis was performed with PANGEA using the gene set “Direct GO Biological Processes” (80). Genes passing the CPM expression filter served as the background gene set for enrichment analysis.

Because the outcrossed wild-type and *cofilin^+/-^* libraries were sequenced on different NovaSeq instruments, potential sequencing-run-associated variation was evaluated by multidimensional scaling (MDS) analysis of TMM-normalized RNA-seq count data from the 18°C samples. The analysis included six outcrossed wild-type samples, six *cofilin^+/-^* samples, and four independent inbred Ore-R controls, with two Ore-R controls represented in each sequencing run. Outcrossed wild-type and *cofilin^+/-^* samples separated primarily along the second leading logFC dimension, whereas Ore-R controls from the two sequencing runs separated mainly along a different dimension (S4 Fig), suggesting that sequencing run was not the primary driver of the genotype-dependent separation. Therefore, sequencing run was not included in the final differential expression analysis.

#### RNA-seq data visualization

RNA-seq data visualizations were generated in R. Volcano plots were generated using ggplot2, with gene labels positioned using ggrepel. MDS coordinates were calculated from TMM-normalized count data using edgeR::plotMDS and visualized using ggplot2. Alluvial plots were generated using ggalluvial and ggplot2. For each alluvial plot, genes from the two indicated differential-expression comparisons were joined by FlyBase ID using the union of genes present in either comparison and classified as Upregulated, Downregulated, Not significant, or Absent from a comparison. Genes classified as Not significant in both comparisons were excluded, and the remaining transitions were summarized by gene count. UpSet plots were generated using the ComplexHeatmap package from FlyBase IDs in the significant up- and downregulated gene sets for the indicated differential-expression comparisons.

#### qPCR analysis and RNA-seq validation

Relative gene expression values for wild-type embryos for *Hsp23*, *Hsp26*, *Hsp68*, *Hsp70, Hsp83*, *Hsc70-4* and *DnaJ-1* were calculated as 2^-ΔΔCτ^ values from qPCR results, as described in Livak and Schmittgen, 2001 and Schmittgen and Livak, 2008 (148, 149). Specifically, Cτ values for individual experimental replicates were normalized by subtracting the corresponding reference gene *robl* Cτ value to obtain ΔCτ. ΔΔCτ values for each experimental replicate were obtained by subtracting each replicate’s ΔCτ value from the mean ΔCτ control value, which was averaged from the three experimental replicates for the 18°C non-stressed wild-type control sample.

For comparison and validation of RNA-seq data with qPCR data, as represented in S2B Fig and S2 Table, the average fold change of counts per million (CPM) of RNA abundance was compared to the average relative gene expression value (-ΔΔCτ) from qPCR for each HSP gene for acute 32°C and chronic 32°C heat-stressed wild-type embryos. Average CPM fold changes for each gene were calculated by dividing the average CPM value for either acute 32°C or chronic 32°C by the average CPM value for 18°C non-stressed wild-type embryos. Data was plotted in MATLAB. Data were fit using a first-order linear regression model, defined as f(x) = p_1_x + p_2_ with slope (p_1_): 0.84 ± 0.22 and intercept (p_2_): 0.66 ± 0.85. Model parameters were estimated by least-squares fitting. The correlation coefficient is *R* = 0.79. Two additional and independent analyses methods yielded similar plots to S2B Fig with adjusted *R* values > 0.78 for all best fit curves.

#### Mass spectrometry analysis

MS2 fragmentation spectra were searched with Proteome Discoverer SEQUEST (version 2.5, Thermo Fisher Scientific) against *in silico* tryptic digested Uniprot *Drosophila melanogaster* (UP000000803). The maximum missed cleavages were set to 2. Dynamic modifications were set to oxidation on methionine (M, +15.995 Da) and protein N-terminal acetylation (+42.011 Da). Carbamidomethylation on cysteine residues (C, +57.021 Da) was set as a fixed modification. The maximum parental mass error was set to 10 ppm, and the MS2 mass tolerance was set to 0.6 Da. The false discovery threshold was set strictly to 0.01 using the Percolator Node validated by q-value. The relative abundance of parental peptides was calculated by integration of the area under the curve of the MS1 peaks using the Minora LFQ node. Mass spectrometry proteomics data were deposited to the ProteomeXchange Consortium via the PRIDE partner repository (150) with the dataset identifier PXD058671.

#### Densitometry

Images of processed Western Blots for S1J Fig were acquired at 300 dpi resolution using the iBright CL1000 instrument and imaging software (Thermo Fisher Scientific). Images were inverted and converted to 8-bit in ImageJ/Fiji and quantified in Adobe Photoshop CC using the Rectangular Marquee tool. For each blot, the integrated intensities per band were normalized against the 18°C wild-type b-actin band so results could be related between biological replicates.

Mean Cofilin protein level in *cofilin^+/-^* embryos is based on n = 3 independent biological replicates.

## Key resources table

| REAGENT or RESOURCE | SOURCE | IDENTIFIER |
| --- | --- | --- |
| **Antibodies** |  |  |
| Affinipure Goat Anti-Mouse IgG(H+L) horseradish peroxidase-conjugated secondary antibody | Jackson ImmunoResearch | Cat#115-035-003<br>RRID:AB_10015289 |
| Affinipure Goat Anti-Rabbit IgG(H+L) horseradish peroxidase-conjugated secondary antibody | Jackson ImmunoResearch | Cat#111-035-003<br>RRID:AB_2313567 |
| Anti-β-actin, Mouse monoclonal | Santa Cruz Biotechnology | Cat#sc-47778<br>RRID:AB_626632 |
| Anti-*Dm* Cofilin, Rabbit polyclonal | Figard et al., 2019 (32) | Cat#Sokac_Dm_Cofilin;<br>RRID:AB_3669047 |
| Anti-Peanut, Mouse monoclonal | Developmental Studies Hybridoma Bank (DSHB) | Cat#4C9H4-c;<br>RRID:AB_528429 |
| Goat anti-mouse IgG (H+L) cross-adsorbed secondary antibody, Alexa Fluor 488 | Invitrogen / Thermo Fisher Scientific | Cat#A-11001<br>RRID:AB_2534069 |
| **Chemicals, peptides, and recombinant proteins** |  |  |
| 2-Chloroacetamide (CAA) | MilliporeSigma | Cat#22790 |
| 4-20% Mini-PROTEAN TGX Pre-cast protein gels, 10-well, 30 uL | Bio-Rad | Cat#4561093 |
| 5-Carboxytetramethylrhodamine (5-TAMRA) | Cayman Chemical | Cat#34598 |
| Acetonitrile, Optima, LC/MS Grade, Fisher Chemical (ACN)I | Acros Organics / Fisher Scientific | Cat#A9554 |
| Actin protein, human platelet, non-muscle, rhodamine-conjugated (G-actin<sup>Red</sup>) | Cytoskeleton, Inc. | Cat#APHR-C |
| Ammonium bicarbonate, 99% (NH<sub>4</sub>HCO<sub>3</sub>) | Thermo Fisher Scientific | Cat#393212500 |
| Aqua-Poly/Mount | Polysciences | Cat#18606-20 |
| BLUeye Prestained Protein Ladder | MilliporeSigma | Cat#94964-500UL |
| ECL Western Blotting Substrate | Pierce / Thermo Fisher Scientific | Cat#32209 |
| Formic acid (FA) | Fisher Scientific | Cat#A11750 |
| Halocarbon 27 oil | MilliporeSigma | Cat#H8773-100ML |
| Halocarbon 700 oil | MilliporeSigma | Cat#H8898-100ML |
| Hoechst 33342 | Invitrogen / Thermo Fisher Scientific | Cat#H3570 |
| Molecular Biology Grade Water | Corning | Cat#46-000-CM |
| Protease Inhibitor Mini Tablets, EDTA-free | Pierce / Thermo Fisher Scientific | Cat#A32955 |
| QC Colloidal Coomassie Stain | Bio-Rad | Cat#1610803 |
| SuperScript II Reverse Transcriptase | Invitrogen / Thermo Fisher Scientific | Cat#18064071 |
| TaqMan Fast Advanced Master Mix | Thermo Fisher Scientific | Cat#4444557 |
| Tris(2-carboxyethyl)phosphine hydrochloride (TECP) | MilliporeSigma | Cat#C4706 |
| Trypsin | Thermo Fisher Scientific | Cat#90305 |

| Critical commercial assays |  |  |
| --- | --- | --- |
| BCA Assay | Pierce / Thermo Fisher Scientific | Cat#23235 |
| iScript Select cDNA Synthesis Kit | Bio-Rad | Cat#1708897 |
| KAPA mRNA HyperPrep Kit | Roche | Cat#KK8581 |
| RNeasy Plus Micro Kit | Qiagen | Cat#74034 |
| Deposited data |  |  |
| Raw RNA-seq data | This paper | NIH BioProject#: PRJNA1368972 |
| Raw mass spectrometry data | This paper | PRIDE Project Accession#: PXD058671; Token: dDxylt5Mqa8L |
| Experimental models: Organisms/strains |  |  |
| Oregon R | Sokac Lab | Available upon request |
| *tsr<sup>1</sup>/CyO* | Bloomington Drosophila Stock Center | Cat#9107; RRID:BDSC_9107 |
| *tsr<sup>N121</sup>/CyO* | Bloomington Drosophila Stock Center | Cat#9109; RRID:BDSC_9109 |
| *white<sup>1118</sup> (w<sup>1118</sup>)* | Gift of Rachel Smith-Bolton Lab | Available upon request |
| Oligonucleotides |  |  |
| LGC BioSearch Technologies Stellaris *twist* Custom RNA FISH probe set, conjugated to TAMRA dye | LGC BioSearch Technologies | This paper; Sequences available upon request |
| Predesigned TaqMan Gene Expression Assay for *Hsp83* | Thermo Fisher Scientific | Cat#4448892 Assay ID: Dm02362342 |
| Custom TaqMan Gene Expression Assay for *Hsp70* | Thermo Fisher Scientific | Cat#4331348, Assay ID: APWC647 |
| Predesigned TaqMan Gene Expression Assay for *Hsp68* | Thermo Fisher Scientific | Cat#4448892 Assay ID: Dm02151262_s1 |
| Predesigned TaqMan Gene Expression Assay for *Hsp26* | Thermo Fisher Scientific | Cat#4448892 Assay ID: Dm01822452_s1 |
| Predesigned TaqMan Gene Expression Assay for *Hsp23* | Thermo Fisher Scientific | Cat#4448892 Assay ID: Dm01822473_s1 |
| Predesigned TaqMan Gene Expression Assay for *DNAJ-1* | Thermo Fisher Scientific | Cat#4448892 Assay ID: Dm01832926_s1 |
| Predesigned TaqMan Gene Expression Assay for *robl* | Thermo Fisher Scientific | Cat#4448892 Assay ID: Dm01821522_g1 |
| Predesigned TaqMan Gene Expression Assay for *Hsc70-4* | Thermo Fisher Scientific | Cat#4448892 Assay ID: Dm02153823_s1 |
| Predesigned TaqMan Gene Expression Assay for *CG15894* | Thermo Fisher Scientific | Cat#4448892 Assay ID: Dm01833202_g1 |
| Predesigned TaqMan Gene Expression Assay for *CG6180* | Thermo Fisher Scientific | Cat#4448892 Assay ID: Dm01790866_s1 |
| Predesigned TaqMan Gene Expression Assay for *CG11034* | Thermo Fisher Scientific | Cat#4448892<br>Assay ID:<br>Dm01804861_g1 |
| Predesigned TaqMan Gene Expression Assay for *CG6282* | Thermo Fisher Scientific | Cat#4448892<br>Assay ID:<br>Dm01841146_m1 |
| Predesigned TaqMan Gene Expression Assay for *tsr* | Thermo Fisher Scientific | Cat#4448892<br>Assay ID:<br>Dm01842418_g1 |
| Software and algorithms |  |  |
| Adobe Illustrator CC | Adobe | https://www.adobe.com/creativecloud.html |
| Adobe Photoshop CC | Adobe | https://www.adobe.com/creativecloud.html |
| IBM SPSS Statistics 29.0.2.0 | International Business Machines Corporation | https://www.ibm.com/products/spss-statistics |
| ImageJ / FIJI | NIH | https://fiji.sc |
| MATLAB | Mathworks | https://www.mathworks.com/products/matlab.html |
| R | R Core Team | https://www.R-project.org/ |
| Other |  |  |
| Aurora UHPLC Column 25 cm×75 µm, 1.7 µm C18 | Ion Opticks | Cat#AUR3-25075C18-TS |
| Cover glass, 24x50 mm, 1 ½ mm thickness | Corning | Cat#2980-245 |
| Fly food R | LabExpress | https://lab-express.com/flyfoodsupplies.htm |
| Heated incubator | PHC Corporation of America | Cat#MIR-154-PA |
| Objective heater and temperature controller | Okolab | Cat#H401-T-Penny |
| Offset flat-tip forceps | Thermo Fisher Scientific | Cat#16-100-116 |
| Snap cap low retention microcentrifuge tubes | Thermo Fisher Scientific | Cat#3448 |
| ZipTip pipette tips | MilliporeSigma | Cat#ZTC18S096 |

## Acknowledgments

We thank Sokac Lab alumni, Liuliu Zheng and Lauren Figard, for laying the foundation for this work. We thank Shrunali Amin for assisting with survival assays. We appreciate discussions regarding stress response with Brian Freeman (Cell and Developmental Biology, University of Illinois Urbana-Champaign). We are grateful for guidance on RNA-seq experiments and analysis from Alvaro Hernandez (DNA Services Lab, Roy Carver Biotechnology Center, University of Illinois Urbana-Champaign); Jenny Drnevich, Jessica Kirkpatrick-Holmes, and Lindsay Clark (HPCBio, Roy Carver Biotechnology Center, University of Illinois Urbana-Champaign); and Kevin Van Bortle (Cell and Developmental Biology, University of Illinois Urbana-Champaign). This work was supported by grants from the National Institutes of Health (R35 GM136384 to A.M.S.; R01 HL126845 to A.K.) and the Chan Zuckerberg Biohub-Chicago to A.K.. SN was supported by Michael A. Recny Fellowship in Biochemistry and Dissertation Completion Fellowship from the Graduate College, University of Illinois Urbana-Champaign. F.R. was supported by a US-Pakistan Knowledge Corridor Scholarship from the Higher Education Commission of Pakistan. The Proteome Exploration Laboratory (California Institute of Technology) is partially funded by Beckman Institute Endowment Funds. Fly stocks were obtained from the Bloomington *Drosophila* Stock Center (supported by NIH grant P40OD018537 and housed at Indiana University, Bloomington). Peanut antibody was obtained from the Developmental Studies Hybridoma Bank (created by the NIH - National Institute of Child Health and Human Development and housed at the University of Iowa, Iowa City).

## Author contributions

All authors contributed to experimental design. N.B., F.R. and D.B. performed fly genetics. N.B. conducted imaging and image analysis. D.B. conducted survival assays. F.R. prepared samples for RNA-seq and conducted qPCR validations. S.N. and A.K. performed RNA-seq analysis. N.B. prepared protein lysates and T.W. and T.C. performed mass spectrometry analysis. N.B., S.N., F.R. and A.M.S. generated figures and tables. N.B., S.N., and A.M.S. wrote the manuscript.

## Supporting information and captions

**S1 Fig.**
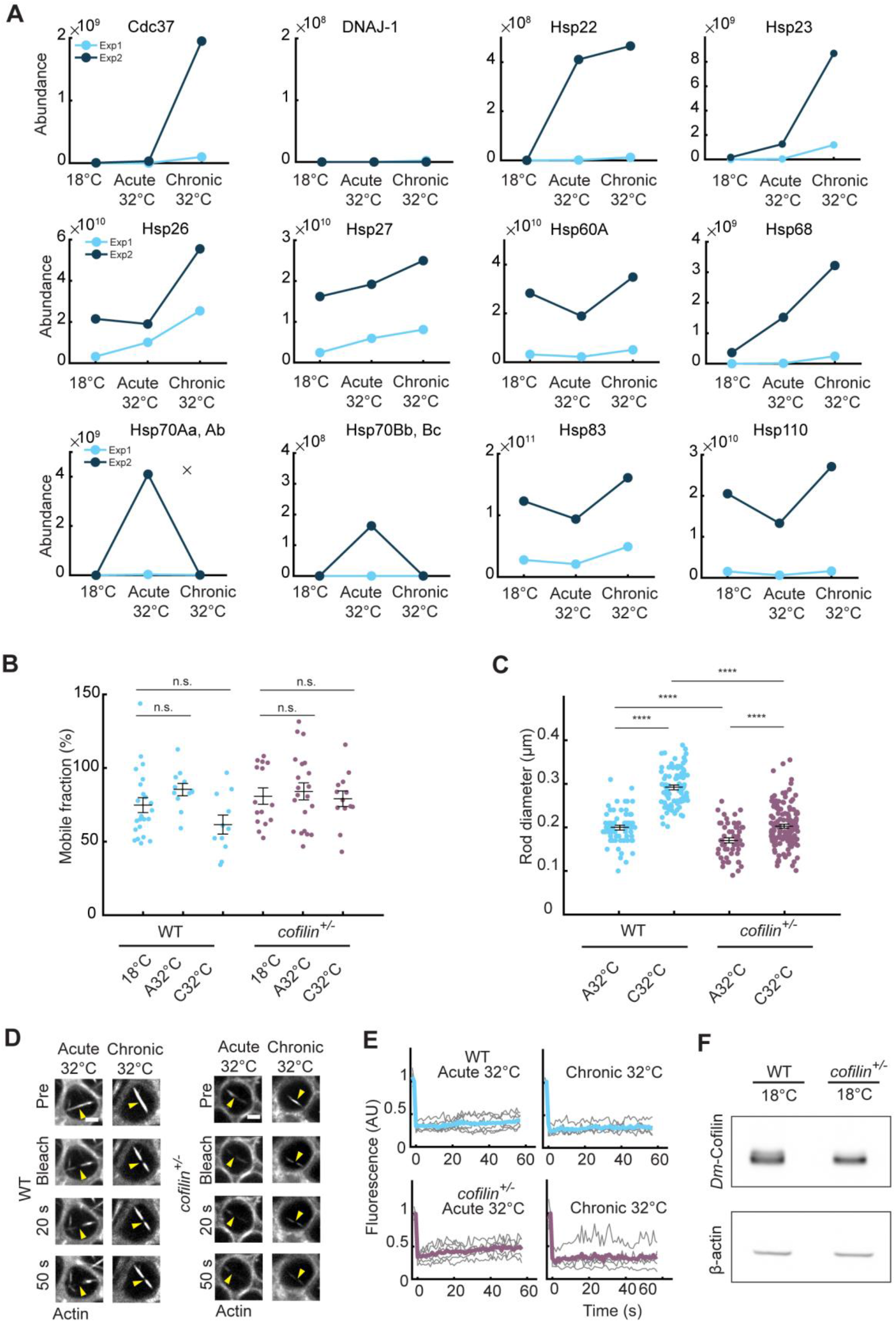
Heat-stressed embryos show induction of HSPs and Cofilin-dependent disruption of cytoplasmic and nuclear actin. Related to Fig 1; S1 Table. (A) Line plots showing raw protein abundance values for HSPs identified from two mass spectrometry experiments for wild-type embryos at indicated conditions. Experiment 1 and 2 shown in light and dark shades, respectively. Data was acquired on an Orbitrap Eclipse Tribrid mass spectrophotometer from experiments repeated for two biological replicates. (B) Dot plot showing percent mobile fraction after photobleaching furrow tip F-actin for WT (cyan) and *cofilin^+/-^* (purple) embryos at indicated temperatures. Each point represents 1 embryo (n ≥ 11 embryos per condition, with 1 furrow bleached per embryo). (C) Dot plot showing rod diameter for WT (cyan) and *cofilin^+/-^* (purple) embryos at indicated conditions (n ≥ 55 rods measured per condition, from 5 nuclei from ≥ 11 embryos per condition). Data shown in Fig 1F. (D) Surface views showing FRAP for nuclear actin rods (G-actin^Red^) in WT and *cofilin^+/-^* embryos at indicated temperatures. Frame “pre” collected immediately prior to bleaching; frames 20 seconds (s) and 50 s collected after bleaching. Yellow arrows show bleached region. Rods in *cofilin^+/-^* embryos start at a lower intensity and the entire rod bleaches with image acquisition due to lack of incorporation of new monomers. Scale bar: 2 μm. (E) Curves showing mean fluorescence recovery after photobleaching for WT (cyan) and *cofilin^+/-^* (purple) image sequences as in (D). Rods of neither genotype show recovery. Individual embryo data shown in gray (n ≥ 5 embryos per condition). (F) Western blot showing reduced Cofilin protein level in *cofilin^+/-^* embryos. β-actin is a loading control. *The mean, shown as horizontal lines in (B) and (C), was calculated from n = 3 independent biological replicates. Error bars represent standard error of the mean (SE). p-values calculated using Student’s t-test. ****p < 0.0001; p* ≥ *0.05 is not significant (n.s.)*.

**S2 Fig.**
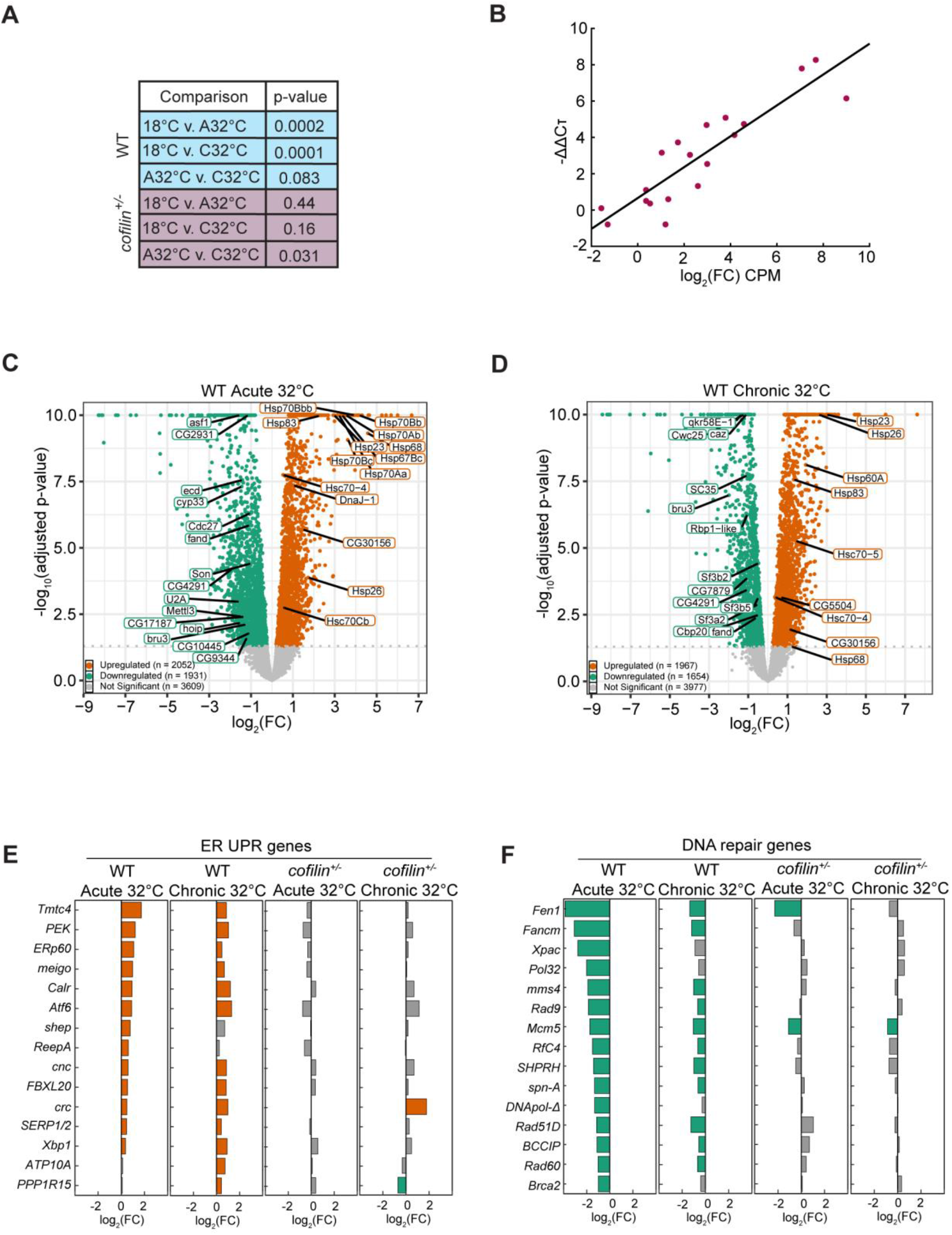
Heat-stressed wild-type embryos show reduced survival and upregulation of HSPs and ER unfolded protein response genes. Related to Fig 2; S2, S3, and S4 Tables. (A) Table showing p-values for within-genotype statistical comparisons of survival for larvae for wild-type (WT; cyan) and *cofilin^+/-^*(purple) embryos at indicated conditions (A, acute; C, chronic; vs., versus). (B) Scatter plot showing mean -ΔΔCτ values from qPCR versus log_2_ of the mean CPM fold change (FC) values from RNA-seq for 7 HSP genes from acute 32°C or chronic 32°C heat-stressed wild-type embryos compared to the 18°C non-stressed condition. Mean CPM FC values were based on n = 6 biological replicates for 18°C WT and acute 32°C WT and n = 5 for chronic 32°C WT; Mean -ΔΔCt values based on n = 3 biological replicates. The correlation coefficient is *R* = 0.79. (C, D) Volcano plots showing -log_10_(adjusted p-value) versus log_2_(FC) for upregulated (orange) and downregulated (green) genes in acute 32°C (C) or chronic 32°C (D) heat-stressed WT embryos versus 18°C non-stressed WT embryos. Number of genes up- or down-regulated, or not significant shown in key. (E and F). Bar plots showing log_2_(FC) values for upregulated (orange) or downregulated (green) genes from highly significant ontologies related to: ER unfolded protein response (E); or DNA repair (F) for WT or *cofilin^+/-^* embryos at indicated conditions. Genes with no significant change in expression shown in gray. Abbreviation in (E) is: unfolded protein response (UPR). *For (A), the mean and standard error of the mean data are shown in Fig 2A. For each condition the mean was calculated from n = 600 embryos total from 3 independent replicates of 200 embryos per replicate. p-values calculated using χ^2^ contingency analysis*. *For (C, D), select HSPs and splicing genes labeled for the up- or down-regulated genes, respectively. Genes associated with the processes of protein folding or splicing chosen due to their common direction of expression observed for most comparisons performed*. *Differential gene expression data in (C-F) based on n = 6 biological replicates for 18°C WT and acute 32°C WT or n = 5 for chronic 32°C WT; and n = 6 for all conditions for cofilin^+/-^*. *Significance value cut-off for differentially expressed genes was FDR < 0.05; and significant ontologies p < 0.01*.

**S3 Fig.**
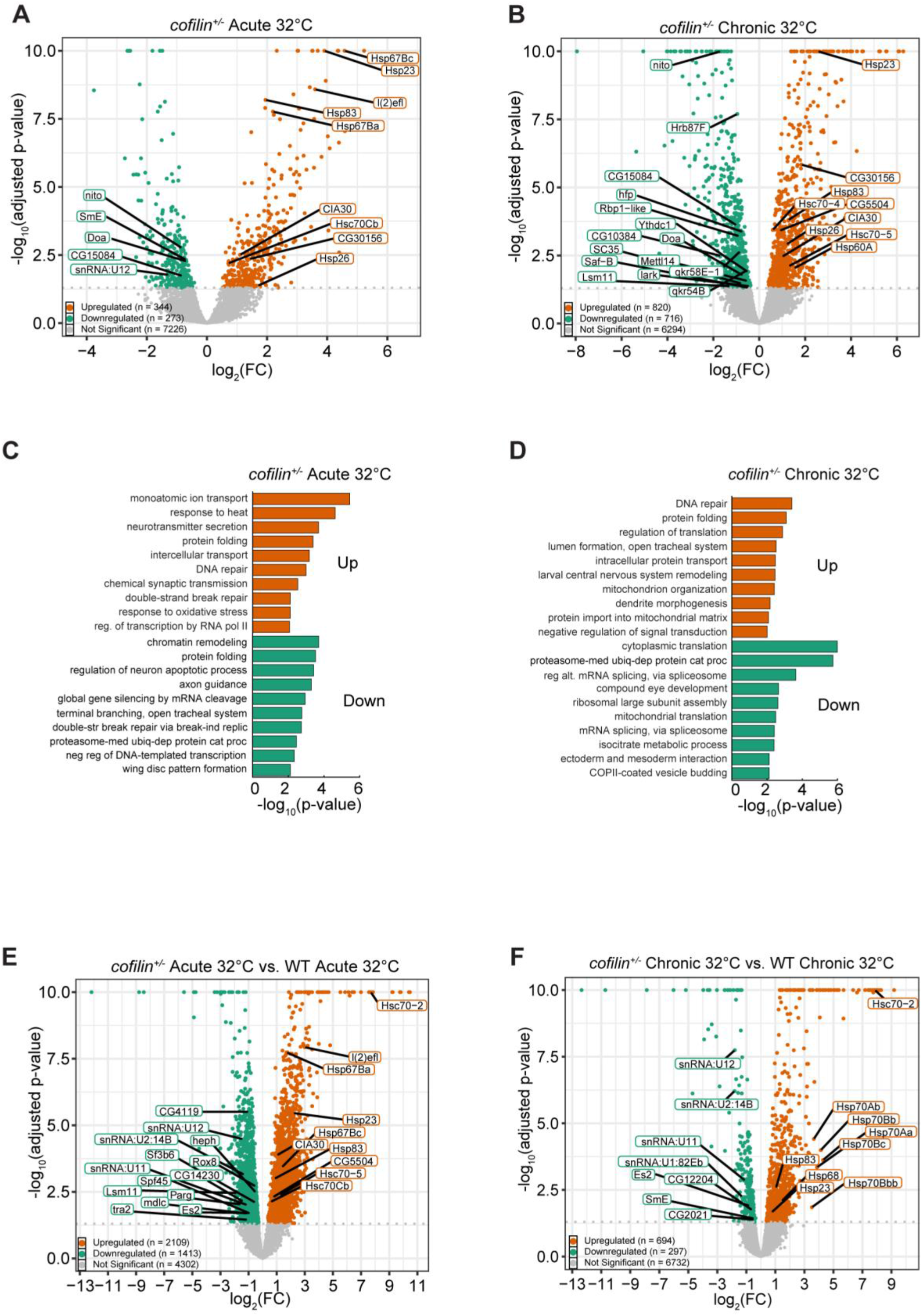
Heat-stressed *cofilin^+/-^* embryos show a distinct stress response compared to wild type. Related to Fig 3; S3, and S4 Tables. (A, B) Volcano plots showing -log_10_(adjusted p-value) versus log_2_(FC) for upregulated (orange) and downregulated (green) genes in acute 32°C (A) or chronic 32°C (B) heat-stressed *cofilin^+/-^* embryos versus 18°C non-stressed *cofilin^+/-^* embryos. Number of genes up- or down-regulated, or not significant shown in key. (C, D) Bar plots showing selected highly significant GO Biological Processes versus -log_10_(p-value) for upregulated (orange) or downregulated (green) gene sets for acute 32°C (C) or chronic 32°C (D) heat-stressed *cofilin^+/-^* embryos versus 18°C non-stressed *cofilin^+/-^* embryos. Abbreviated processes are: double-strand break repair via break-induced replication; proteasome-mediated ubiquitin-dependent protein catabolic process; negative regulation of DNA-templated transcription; regulation of alternative mRNA splicing, via spliceosome. (E, F) Volcano plots showing -log_10_(adjusted p-value) versus log_2_(FC) for upregulated (orange) and downregulated (green) genes in the *cofilin^+/-^*acute 32°C versus (vs.) WT acute 32°C comparison (E) or the *cofilin^+/-^* chronic 32°C vs. WT chronic 32°C comparison (F). Number of genes up- or down-regulated, or not significant shown in key. *For (A, B, E and F), select HSPs and splicing genes labeled for the up- or down-regulated genes, respectively. Genes associated with the processes of protein folding or splicing chosen due to their common direction of expression observed for most comparisons performed*. *Differential gene expression data in (A-F) based on n = 6 biological replicates for 18°C WT and acute 32°C WT or n = 5 for chronic 32°C WT; and n = 6 for all conditions for cofilin^+/-^*. *Significance value cut-off for differentially expressed genes was FDR < 0.05; and significant ontologies p < 0.01*.

**S4 Fig.**
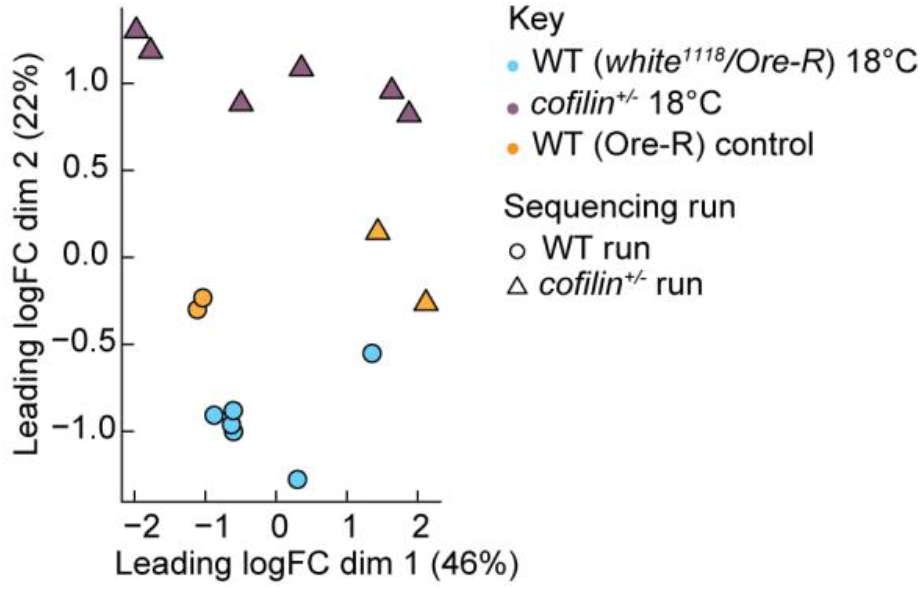
Assessment of sequencing-run-associated variation in RNA-seq samples. Related to Fig 4. Multidimensional scaling (MDS) plot of TMM-normalized RNA-seq count data from non-stressed 18°C outcrossed WT and *cofilin^+/-^*embryos together with independent inbred Ore-R controls represented across the two sequencing runs. Each point represents one biological replicate. Colors indicate genotype/control group and shapes indicate sequencing run. The analysis includes n = 6 WT, n = 6 *cofilin^+/-^*, and n = 4 Ore-R controls, with two Ore-R controls represented in each sequencing run.

**S1 Table. Mass spectrometry data for HSPs.**

Excel file containing mass spectrometry data.

**S2 Table. Validation of RNA-seq data by qPCR.**

Excel file containing qPCR and CPM data.

**S3 Table. Differentially expressed genes from RNA-seq.**

Excel file containing differentially expressed genes from RNA-seq analysis.

**S4 Table. Enrichment analysis for differentially expressed genes identified by RNA-seq.**

Excel file containing gene ontology analysis (GO) data.

